# C-COMPASS: A Neural Network Tool for Multi-Omic Classification of Cell Compartments

**DOI:** 10.1101/2024.08.05.606647

**Authors:** Daniel Thomas Haas, Eva-Maria Trautmann, Xia Mao, Mathias J. Gerl, Christian Klose, Xiping Cheng, Jan Hasenauer, Natalie Krahmer

## Abstract

Functional compartmentalization in eukaryotic cells is essential for maintaining physiological processes. The development of systematic organelle proteomic techniques, such as Protein Correlation Profiling (PCP) and Localization of Organelle Proteins by Isotope Tagging (LOPIT), has enhanced our understanding of organelle dynamics and protein localization. However, the complexity of the data and the need for advanced computational skills limit their accessibility. To address this, we introduce C-COMPASS, an open-source software featuring a user-friendly interface that utilizes a neural network-based regression model to predict the spatial distribution of proteins across cellular compartments. C-COMPASS manages complex multilocalization patterns and integrates protein abundance information to model organelle composition changes under various biological conditions. Using C-COMPASS, we mapped the organelle proteomic landscapes of humanized liver mice in different metabolic states and modeled changes in organelle composition to provide insights into cellular adaptations. Additionally, we extended cellular maps to the lipid level by co-generating protein and lipid profiles. C-COMPASS trains neural networks with marker protein profiles to predict lipid localizations, enabling parallel mapping of lipid and protein localization. This approach overcomes the lack of knowledge of organelle-specific lipid markers and identifies previously unknown organelle-specific lipid species. C-COMPASS offers a comprehensive solution for studying organelle dynamics at the multi-omics level, designed to be accessible without requiring extensive computational expertise or specialized high-performance computing equipment.

## Main

Cellular compartmentalization is a fundamental principle of eukaryotic cell organization, creating distinct functional modules that facilitate biochemical reactions, segregate toxic metabolites, and enable precise regulation of signal transduction and intracellular transport^1,2^. This compartmentalization is tailored to the unique functions of different cell types and plays a crucial role in cellular differentiation and specialization^3,4^. Understanding compartmental changes is essential for unraveling cellular mechanisms and dysfunctions, providing insights into cell biology and disease development.

Organelles, which are characterized by their unique protein and lipid composition, are dynamic and rapidly adapt to environmental changes. The interaction and physical attachment of organelles render complete purity of organelle isolation difficult to achieve. This is further compounded by high-sensitivity proteomics, which can detect significant portions of the proteome, even in highly pure organelle preparations^5^. Advanced organelle proteome profiling techniques have partially overcome these limitations. These include Protein Correlation Profiling (PCP)^6,7^, Localization of Organelle Proteins by Isotope Tagging (LOPIT)^8^, SubCellBarCode^9^, and Dynamic Organellar Maps (DOMs)^10^. These methods use differential or density-gradient centrifugation to separate cellular components by size and density and liquid chromatography-tandem mass spectrometry (LC-MS/MS) to quantify protein abundance across these fractions (Fig.1a-d). This creates profiles indicative of organelle-specific protein distribution, aiding in distinguishing genuine compartmental proteins from contaminants (Fig.1e). These methods provide localization data for thousands of proteins across multiple compartments in a single experiment, enhancing our understanding of intracellular protein dynamics and functions^11–13^.

Streamlined cellular fractionation workflows combined with high-sensitivity, high-throughput LC-MS/MS facilitate mapping of protein localization and dynamics across biological conditions. However, the resulting datasets are complex and contain abundance profiles for thousands of proteins, with an estimated 50-60% of proteins having multiple localizations^14^, making organelle classification challenging. Computational methods, including Bayesian models and supervised machine learning, such as support vector machines (SVMs)^12,15^, leverage organelle marker proteins to classify others into specific compartments. However, no existing techniques effectively address the challenges of handling proteins with multiple localizations and integrating organelle and protein abundance data, which are additional factors that determine organelle composition. Moreover, their implementation typically requires coding skills, which presents a limitation for users lacking computational expertise.

Here, we introduce C-COMPASS (Cellular COMPartmentclASSifier), a standalone open-source software, providing a reproducible analysis pipeline in a graphical user interface. C-COMPASS features neural-network-based regression models that predict proteome-wide spatial distributions across multiple compartments from cellular and tissue fractionation data. C-COMPASS advances upon earlier methods by addressing complex multilocalization patterns of proteins, moving beyond single-compartment predictions. Previous SVM workflows, which were trained with profiles of known marker proteins, were limited to predicting a single organelle^8,10,12,15,16^ (Fig. 1f), leading to issues in classification reliability for proteins with multiple localization. C-COMPASS addresses this limitation using a neural network-based regression model that predicts the proportional localization of each protein across multiple organelles to account for the complex nature of cellular compartmentalization. Furthermore, C-COMPASS integrates protein and organelle abundance derived from co-analyzed whole-cell or tissue proteomes to estimate compositional changes of cellular compartments across biological conditions and effectively model organelle composition. Using our tool, we created cellular maps of mice with humanized livers in various metabolic states, mapping protein relocalization events and organelle reprogramming.

**Figure 1.**
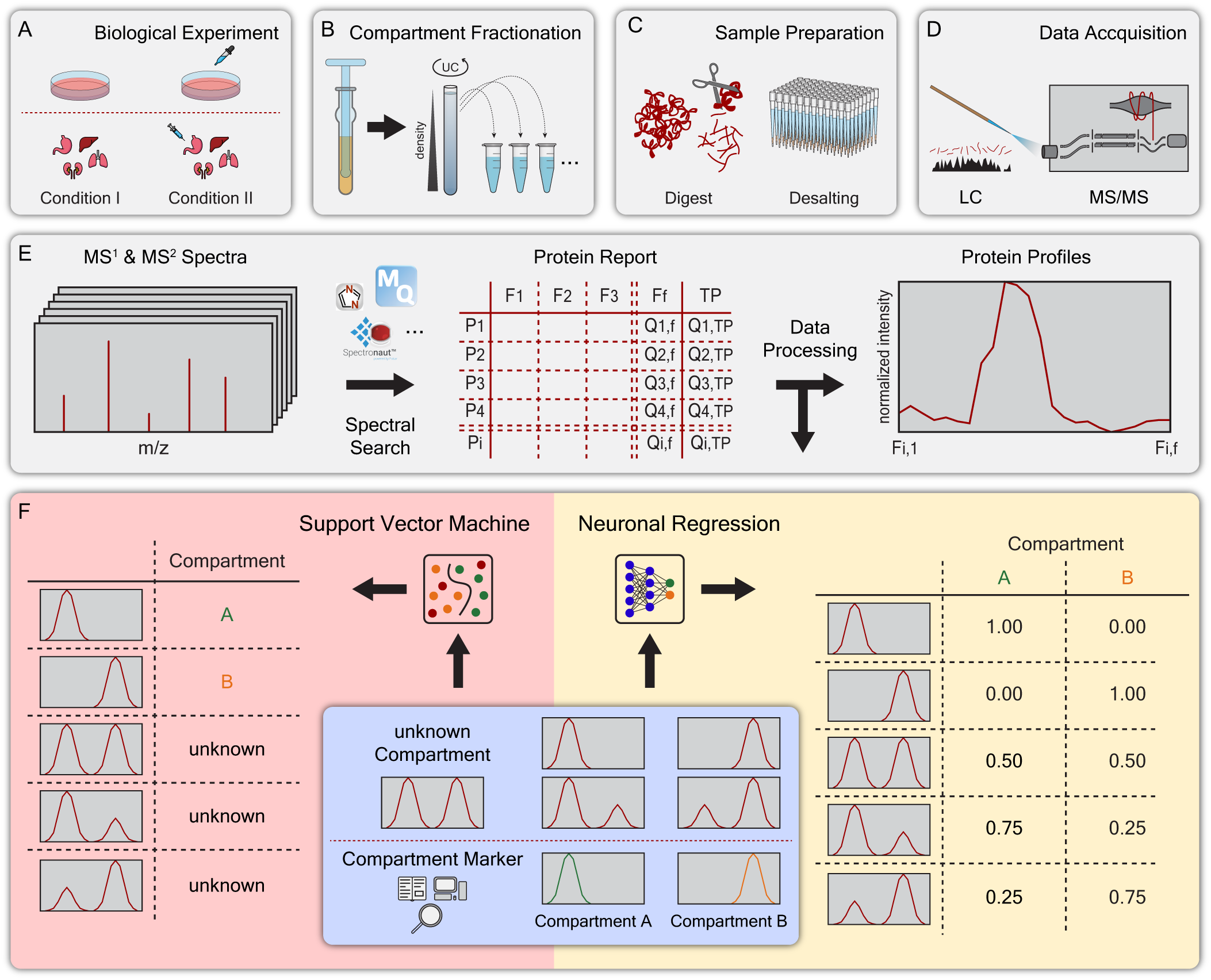
Overview of workflow to generate organellar maps. (A) Cells or tissue samples are collected from the biological conditions of interest in biological replicates. (B) Samples are lysed while keeping organelles intact, and compartments are separated, for example, based on density centrifugation. (C)-(D) Compartment fractions are prepared for LC-MS/MS, and proteins are quantified. (E) Abundance information for proteins (P1-Pi) in all organelle fractions (F1-Ff) and total proteome information (TP) is obtained via standard LC-MS/MS workflows. Quantitative information from pivot report tables is used to generate abundance profiles by scaling to an area of 1 or by MinMax scaling across the fractions. (F) ML-based localization predictions require marker proteins specific to single compartments. Traditional methods like SVM produce binary predictions, struggling with multiply localized proteins, whereas neuronal regression models assess contributions from multiple compartments to a protein profile.

Additionally, we demonstrate the versatility of C-COMPASS beyond proteomics. By co-training neural networks with concurrently generated proteome, C-COMPASS enables the prediction of subcellular localization of in parallel generated lipidomic profiles, identifying organelle-specific lipid species. This capability allows for the first time the prediction of lipid compositions of organelles in a systematic manner. Overall, C-COMPASS provides a comprehensive deep-learning tool for mapping cellular organelles in healthy and diseased states without requiring programming skills.

## Results

### Computational Workflow

C-COMPASS analysis is based on experimentally generated protein abundance profiles, independent of the LC-MS/MS acquisition method, such as data-dependent acquisition (DDA) or data-independent acquisition (DIA). It can incorporate various labeling approaches, including label-free, Tandem Mass Tag (TMT), or Stable Isotope Labeling by Amino acids in Cell culture (SILAC). To annotate protein localizations based on cellular fractionation data, C-COMPASS requires two inputs: (1) a pivot report table listing identified proteins and their intensities across fractions, and (2) a marker set of proteins typically localized to single compartments. C-COMPASS provides predefined marker lists derived from published sources and experimentally generated databases, such as Protein Atlas or Mapofthecell, and users can also import custom lists from tabular files.

Upon uploading data via the user interface and defining biological conditions, replicate numbers, and fractions, C-COMPASS normalizes each protein profile within replicates and combines them into a unified dataset, which is filtered based on occurrence across replicates (Fig. 2a). The framework then matches organelle annotations of markers with the dataset. To address variability in marker protein availability, we implemented upsampling by combining three randomly selected profiles from each underrepresented batch into artificial profiles with added random noise. Additionally, we simulate multiple localizations by mixing marker profiles from two compartments for a subset of the upsampled batch (Fig. 2a).

**Figure 2.**
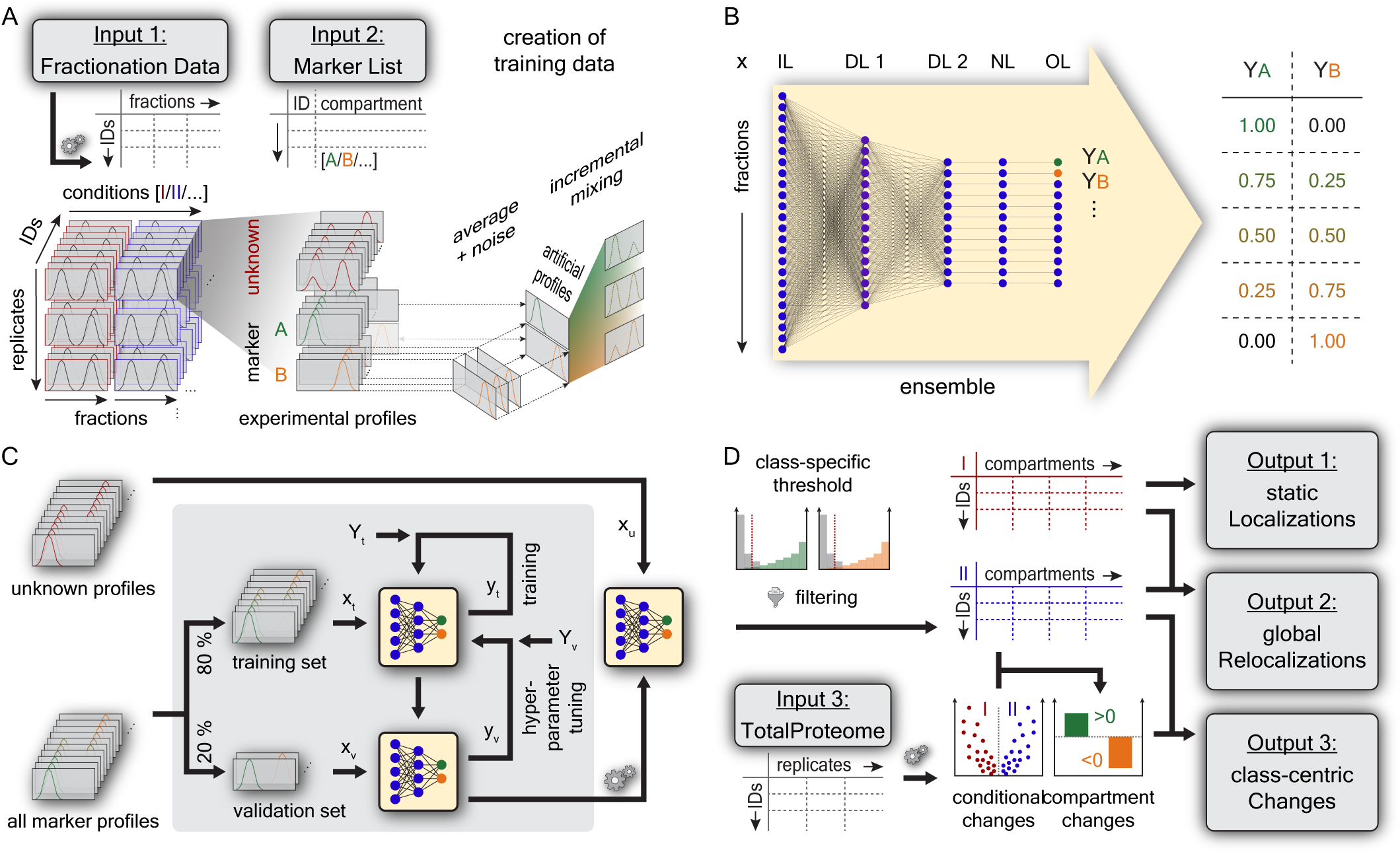
The C-COMPASS framework. (A) Fractionation data is imported into C-COMPASS (Input 1) and filtered for valid values. Profiles are scaled to an area of 1, creating a data matrix with valid profiles across replicates and conditions (red and blue). These protein lists are matched with a marker list (Input 2) from literature, containing compatible identifiers and main compartment assignments. To balance marker batch sizes, artificial profiles are created by adding random noise to randomly selected profiles from underrepresented classes. Marker profiles from two compartments are incrementally mixed to represent dual localizations. (B) The input layer (IL) of the neural network matches the number of fractions in a replicate. Two hidden dense layers follow: the first layer (DL1) size is optimized via hyperparameter tuning, and the second layer (DL2) size corresponds to the number of compartments. A normalization layer (NL) ensures neuron data sums to 1, and the output layer (OL) provides the network output (Y) for each compartment. (C) The network topology is optimized through hyperparameter tuning. Marker profiles are split into a training set to compare actual values (Y_t_) with network output (y_t_) and a validation set to evaluate performance on unseen data by comparing actual values (Y_v_) with network output (y_v_). The best topology is used for processing the entire dataset, including marker and unknown profiles. (D) Neural network output is filtered by setting a threshold to cover 95% of false positive values per compartment, setting all values below the threshold to 0, and re-normalized to sum to 1. The results provide CC values for all compartments (Output 1), which can be compared across biological conditions to estimate relocalizations (Output 2). Total proteome values (Input 3) can be used to assess changes in protein expression levels and compartment abundance, normalizing the relocalization information to generate class-centric statistics (Output 3).

C-COMPASS uses a neural network-based regression model trained on experimental and artificially generated profiles (Fig. 2b). The network has five layers, with input layer (IL) size based on the number of fractions to fit the length of profiles as input values (x). Two dense layers (DL1 and DL2) follow, with DL2 matching the number of compartments. The size of DL1 is a tunable hyperparameter. A normalization layer (NL) adjusts contributions to sum to one, leading to the final output layer (OL). The learning rate of the model and optimizer are tuned for each run. The network is trained multiple times using the marker list. The output is a matrix of prediction values per compartment, aggregated to derive a final class contribution (Y) for each compartment per condition. Hyperparameter tuning is performed by splitting marker profiles into a training set (80%) and a validation set (20%), allowing for continuous validation and optimization of the network performance (Fig. 2c). During training, the network learns by comparing output values (y_t_) with actual compartment annotations (Y_t_). The model is then tested on the validation set, refining hyperparameters based on output comparisons (y_v_ vs. Y_v_). This iterative process identifies the best network architecture. Finally, the optimal model is trained on all marker profiles, enabling it to predict class contributions for the entire protein list.

The output values for marker proteins are analyzed by compartment, categorizing them into specific compartment markers and markers for all other compartments (Fig. 2d). Since the prediction is not binary, ranges of values are classified as true or false positives. A threshold is set where 95% of false positives fall below this value; neural network output values below this threshold are set to zero. The remaining values are re-normalized so their sum across compartments for each protein equals one. These normalized values, called class contributions (CC), represent each compartment’s contribution to a protein profile, resulting in a data matrix indicating protein distribution across compartments. These static localizations per condition are the first output of C-COMPASS. CC values are used for comparative analyses between conditions. The difference in compartment-specific CC values, called relocalization (RL), ranges from -1 to 1. By using the outputs from running the neural network as an ensemble, we compute a p-value and Cohen’s distance (D) for each compartment. We also generate a relocalization score (RLS) with corresponding p-value, and a distance score (DS). The RLS, ranging from 0 (no relocalization) to 2 (full relocalization), sums absolute RL values across all compartments. The DS, an unlimited positive value, ranks relocalizations by summing the absolute D values. These metrics form the second C-COMPASS output, referred to as global relocalizations. To further investigate biological system changes, we integrate total proteome data. We normalize all relocalization values with the corresponding conditional changes in protein expression levels. Additionally, we combine the previously generated spatial information with total proteome data to estimate changes in compartment abundance. This third C-COMPASS output offers a class-centric perspective on protein level changes within various compartments, addressing systematic changes. We applied and validated the C-COMPASS pipeline using various published and unpublished datasets to confirm the performance of each individual pipeline step.

### Assessing Multiple organelle predictions

To evaluate the performance of C-COMPASS in predicting multi-organelle localization, we applied it to a dataset containing information on the localization of 5,530 proteins in human white adipocytes using PCP to separate compartments across 24 gradient fractions^17^ (Fig. 3a). We compared the SVM output for differentiated adipocytes from the study with the multiple organelle predictions generated by C-COMPASS using the same study-derived marker set, which encompasses 12 distinct compartment classes (Fig. 3b). Initially, we assessed C-COMPASS performance with and without upsampling. The original marker list displayed significant discrepancies in the number of markers across compartments. By applying the upsampling pipeline, we mitigated these discrepancies and compared the original and upsampled datasets using UMAP visualization (Fig. 3c). Post-upsampling, the batches were uniformly distributed across all compartments while preserving distinct clusters. Adding random noise to the profiles resulted in a similar distribution between artificially and experimentally generated protein profiles (Fig. 3c). We then evaluated performance using precision, recall, and F1 score, standard metrics for machine-learning models, and compared predictions with and without upsampling and profile mixing, based on raw neural network outputs (Fig. 3d). For example, lipid droplets (LDs), initially with only 25 markers, showed minimal correct predictions with an average F1 score below 0.01, which increased to 0.50 after upsampling. Conversely, mitochondrial proteins, with 500 markers, performed well even without upsampling, achieving an F1 score of 0.82, which improved to 0.86 post-upsampling. On average, across all twelve compartments, precision increased from 0.44 to 0.60, recall from 0.47 to 0.66, and F1 score from 0.46 to 0.63 after upsampling. Additionally, the results show that mixing profiles for training the network, does not reduce prediction reliability.

**Figure 3.**
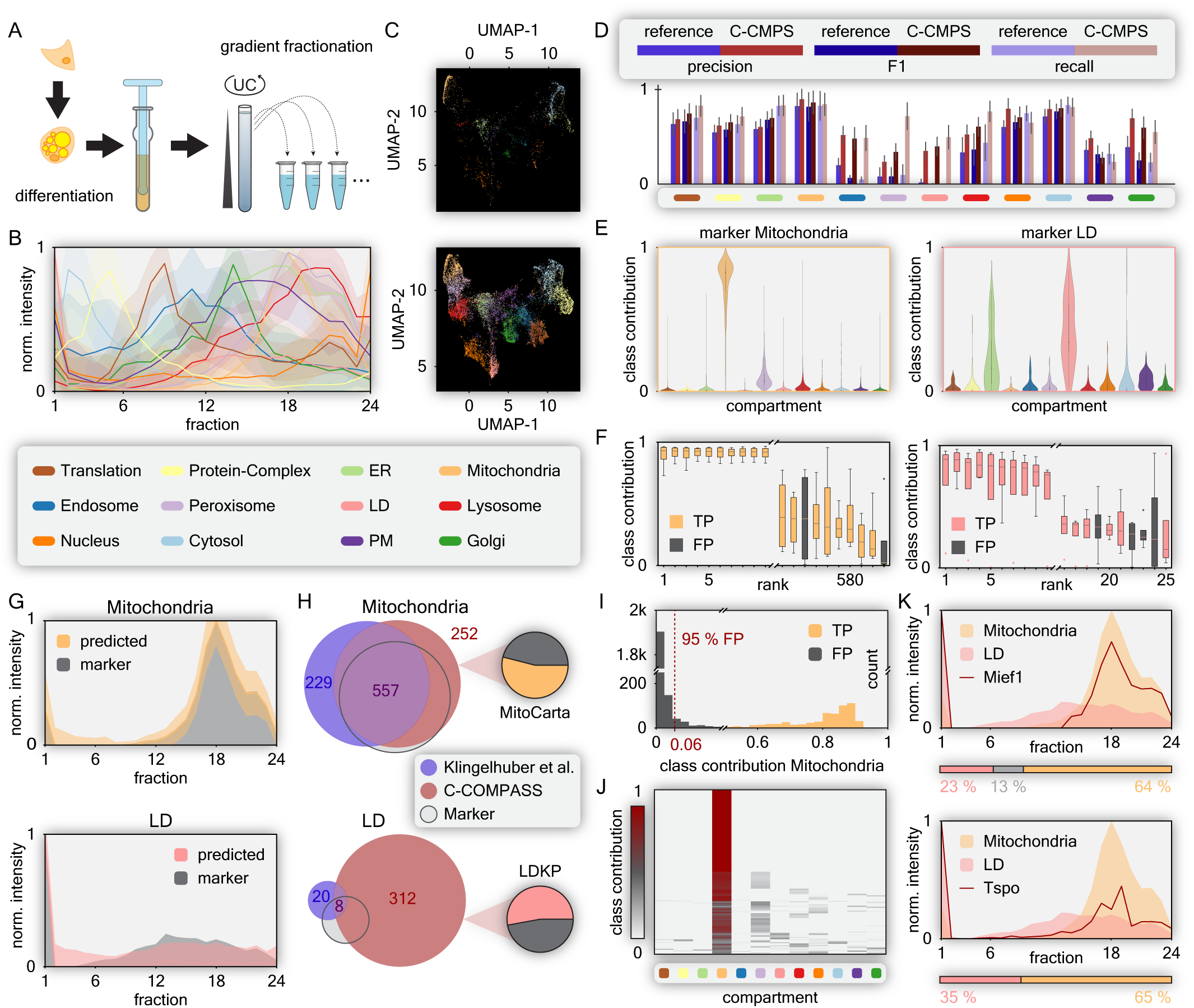
C-COMPASS performance in predicting multiple localizations. (A) Scheme to generate an organellar map of adipocytes via PCP. (B) Profiles of compartment markers (median from 4 biological replicates, error bands represent standard deviations). (C) UMAPs displaying data points across all replicates for compartment markers. UMAPs trained (n_neighbors = 10, n_components = 2, metric = ‘cosine’, min_dist = 0.1, spread = 1.0, random_state = 42, learning_rate = 1.0, n_epochs = none, init = ‘spectral’, and transform_seed = 42) on upsampled dataset (lower) were applied to represent the original marker proteins (upper). (D) Precision, F1, and recall metrics are shown without upsampling (blue) and with C-COMPASS (C-CMPS) upsampling and mixing (red). Proteins were assigned to compartments based on the highest neural network outputs. Error bars for reference values are from network ensemble values, while C-CMPS values include ensemble and multiple upsampling rounds, representing standard deviations. (E) For Mitochondria and LD, raw neural network outputs from all neurons derived from marker proteins are summarized. Violins are width-scaled, inner lines represent quartiles, and dots represent median values. (F) Boxplots display ranked network outputs for Mitochondria and LD marker proteins. True positives (TP) are colored, and false positives (FP) are grey. Medians are shown with middle lines, boxes indicate interquartile ranges (IQR), whiskers extend to 1.5 times the IQR, and dots represent outliers. (G) Median profiles for MinMax scaled protein profiles. Marker proteins are shown in grey, with Mitochondria (orange) and LD (red). Error bands indicate standard deviations. (H) Venn diagrams comparing proteins with the highest neural network output values for Mitochondria (upper) and LD (lower) from C-COMPASS (red) with original predictions by Klingelhuber et al.^17^ (blue). Colorless circles show intersections with marker proteins. C-COMPASS unique predictions are compared with MitoCarta^19^ and LDKP^20^. (I) Histogram of neural network output values for Mitochondria. Orange bars represent markers for Mitochondria, grey bars represent other organelles, and the red dashed line shows the filtering threshold. (J) Heatmap of CC values per compartment, filtered for proteins at least partially localized to Mitochondria. (K) Example MinMax scaled protein profiles for two proteins localized to Mitochondria and LD. Colored areas in line plots represent median marker profiles for Mitochondria (orange) and LD (red). Horizontal bars indicate compartment distribution as predicted by C-COMPASS.

To provide an additional performance evaluation, we assessed the pre-filtered distribution of class predictions for marker proteins across the different compartments. These marker proteins are chosen for their primary, exclusive localization to specific compartments, and they exhibited high prediction values for their designated compartments and low values for others, which is consistent with biological expectations (Extended Figure 1a). For instance, proteins localized to the mitochondria, which typically have a single organelle localization due to their distinctive targeting signals, showed high prediction values for mitochondria and low values for other compartments (Fig. 3e). The ranking of mitochondrial prediction values across all mitochondrial marker proteins revealed only two false positives among the 583 proteins (Fig. 3f). LD proteins, which often have multiple localizations, showed higher values for other compartments. However, the C-COMPASS predictions were consistent with the biological context of the compartments, as proteins are directed to LDs either from the cytosol (CYTOLD) or endoplasmic reticulum (ERTOLD)^18^. This was reflected by LD markers showing high values for LDs and elevated values for the ER and cytosol (Fig. 3e). Ranking LD prediction values across all compartment markers revealed four false positives among the 26 predicted on the LD (Fig. 3f). Overall, the profiles closely matched the marker profiles of their predicted compartments (Fig. 3g). When comparing the C-COMPASS predictions with the SVM-based output from the original study by Klingelhuber et al.^17^ (Fig. 3h), we observed 53.7% overlap in the mitochondrial predictions. C-COMPASS predictions for mitochondria showed greater overlap with marker proteins than SVM-based predictions. Additionally, 54% of the unique findings from C-COMPASS were confirmed as mitochondrial, according to MitoCarta^19^, indicating improved accuracy. For LD predictions, the number of predicted LD proteins increased from 28 in the original study to 320, with 53% confirmed by the LD Knowledge Portal (LDKP)^20^, indicating enhanced identification of LD proteins compared to SVMs (Fig. 3h). C-COMPASS outperformed SVMs in correctly assigning marker proteins to their respective compartments for 11 out of 12 compartments. The only exceptions were plasma membrane (PM) predictions, where SVMs showed better performance (Extended Figure 1b).

When assessing organelle prediction values for mitochondria, we noticed that several marker proteins for other compartments had low mitochondrial prediction values, indicating potential false positives rather than genuine localizations. To address this, we applied a filtering step, setting a threshold to a compartment individual 95% false-positive rate. By implementing this threshold, we successfully filtered out potential false positives (markers for other compartments) without losing true positives, as shown for mitochondrial marker proteins (Fig. 3i). These filtered values, named class contribution (CC), were then used to predict multiple organelle localizations, as demonstrated for the mitochondrial compartment (Fig. 3j), enabling the identification of exclusive organellar proteins and proteins with multiple localizations. Examples of colocalization across mitochondria and LDs include mitochondrial outer membrane proteins such as Mief1 (Fig. 3k), which is involved in fission regulation at organelle contact sites^21^, and the mitochondrial cholesterol transporter Tspo (Fig. 3k). Tspo has been previously described to reside at mitochondrial-nuclear contact sites and to mediate cholesterol redistribution in the cell in a complex with protein kinase A (PKA)^22^.

### Assessment of Handling Versatility in Experimental Workflows and Evaluation of Comparative Analyses

To demonstrate the broad applicability of C-COMPASS across various organelle separation techniques and proteomic labeling strategies and to evaluate its reliability in identifying protein relocalization events across conditions, we applied the pipeline to a published dataset by Mulvey et al.^23^. This dataset utilized the HyperLOPIT approach, based on differential centrifugation and TMT labeling proteomics, to assess changes in protein localization in the human monocytic leukemia cell line THP-1 upon induction of an immune response via treatment with lipopolysaccharide (LPS) (Fig. 4a).

**Figure 4.**
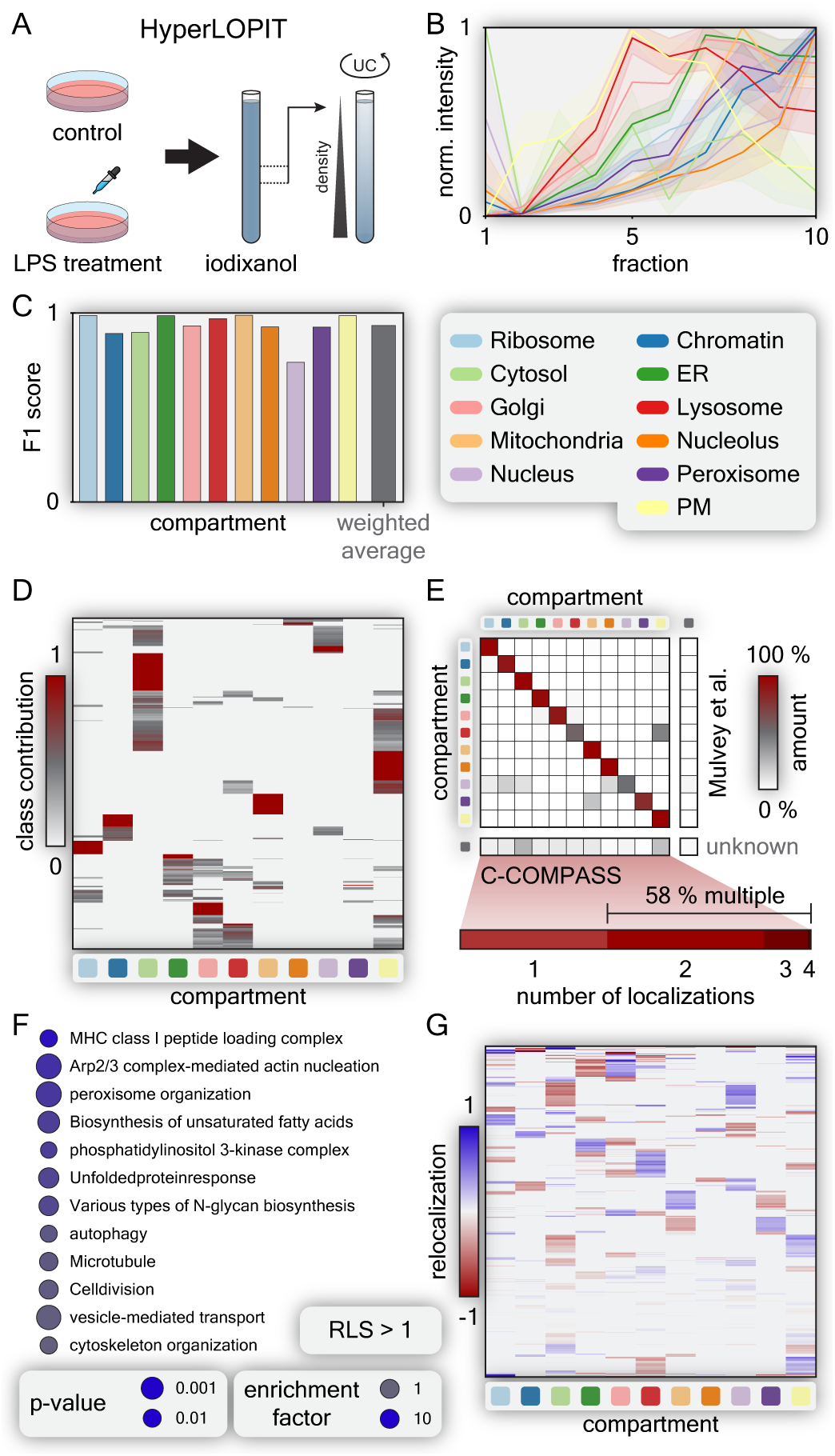
The application of C-COMPASS to different fractionation workflows. (A) Experimental design to generate a cellular map of LPS treatment via HyperLOPIT by Mulvey et al.^23^. (B) Profiles for compartment markers, with line values representing median values derived from median profiles across replicates for control condition, and error bands indicating standard deviations. (C) Bar plot showing F1 scores for each compartment in the control condition. Main organelle assignments were determined using the highest CC value per protein. The grey bar represents the average F1 value weighted by batch sizes of marker proteins. (D) Heatmap of CC values across all compartments in the control condition. (E) Correlation matrix comparing original compartment associations by Mulvey et al.^23^ with C-COMPASS main compartment predictions for the control condition. The lower bar shows the percentage of proteins with varying numbers of localizations, for proteins not assigned in the original study. (F) GO-annotation, KEGG and Keyword enrichment analysis for proteins with RLS >1 upon LPS treatment (one-sided Fisher’s exact test, FDR <0.15) (G) Heatmap of RL values for each compartment across all proteins upon LPS treatment.

Although the reduced number of fractions resulted in a greater overlap in organelle distribution profiles compared to PCP profiles (Fig. 4b, Extended Fig. 2a) the F1 scores achieved by C-COMPASS (0.74 to 0.99 with a weighted average of 0.93 in the control condition, and 0.74 to 1.00 with a weighted average of 0.91 for LPS-treated condition), indicate that the main compartment predictions remain highly reliable (Fig. 4c, Extended Fig. 2b). We found that 49.4% of proteins exhibited multiple localizations, consistent with previously reported numbers from microscopy-based methods^14^ (Extended Fig. 2c). A heatmap of the compartment associations shows the CC values across compartments for these proteins, which is consistent with the biological context (Fig. 4d, Extended Fig. 2d-e). For example, proteins moving through the secretory pathway often localize to the Golgi apparatus, endosomes, and plasma membrane, whereas most mitochondrial proteins targeted via specific signaling peptides are uniquely localized. A comparison matrix between the original predictions by Mulvey et al. and the main C-COMPASS predictions (Fig. 4e, Extended Fig. 2f) revealed a strong correlation. Differences include that for many lysosomal predictions of the original study, C-COMPASS predicted dual lysosomal-plasma membrane localizations and for proteins predicted in the original study as nuclear, we predicted multiple localizations across the nucleus, ribosome, chromatin, and cytosol, potentially reflecting the dynamic nature of proteins that shuttle between these compartments (Fig. 4e). Additionally, many proteins (58%) classified as unknown in the original study exhibited multiple localizations according to C-COMPASS (Fig. 4e), which explains their classification difficulty with traditional machine learning classifiers.

Next, we validated the C-COMPASS approach for identifying protein relocalizations using the dataset. To identify proteins with the strongest changes in localization prediction, we first filtered for a RLS with a threshold of >1, indicating a 50% change in organelle assignments. To identify the most reliable hits, we plotted the C-COMPASS distance metric against the RLS p-value. Higher values indicate outliers that exhibit greater reproducibility across biological replicates (Extended Fig. 2g). Proteins with an RLS >1 showed an enrichment of proteins involved in pathways known to be activated upon LPS treatment, including immune response-related terms such as the MHCI peptide loading complex^24^, the unfolded protein response^25^, actin nucleation^26^, cell division^27^, lipid metabolism^28^, as well as processes highlighted in the original study to be affected, including vesicular transport and cytoskeleton remodeling (Fig. 4f). Additionally, we observed significant reorganization of the Pi3K pathway, known to be activated upon LPS stimulation^29^. In concordance with the original study using the HyperLOPIT workflow, which utilizes a T-Augmented Gaussian Mixture Model with Bayesian computation performed using a Markov Chain Monte Carlo method (TAGM-MCMC) to assign proteins to distinct subcellular regions, quantify the uncertainty in their allocation, and identify relocalizations, we found a high number of translocation events from the lysosome to the plasma membrane and from the cytosol to the nucleus (Fig. 4g). This indicates that C-COMPASS can detect known and previously validated protein relocalizations under the specific treatment condition.

### Organelle composition modeling using reduced fraction PCP

Organelle composition is determined by three main parameters: (I) protein localization, (II) protein abundance, and (III) organelle abundance. To predict changes in organelle composition, C-COMPASS integrates these factors. We applied this enhanced workflow to generate organelle maps from mice with humanized livers, allowing the study of organelle remodeling in human hepatocytes under various metabolic conditions and for the first time in a human liver context.

FRG (Fah/Rag2/Il2rg-deficient) mice, whose fumarylacetoacetate hydrolase (Fah)-deficient hepatocytes die due to the accumulation of toxic tyrosine metabolites, were humanized by engraftment of the liver with FAH-proficient human hepatocytes^30^ carrying a *PNPLA3* mutation known to exacerbate lipid accumulation. Using this established humanized model for metabolic dysfunction-associated fatty liver disease (MASLD)^31^, we compared the cellular architecture of humanized livers in mice fed a standard chow diet or a high-fat, high-fructose (HFHF) diet for five weeks and assessed the reorganization of cellular architecture after an overnight fast in the HFHF-condition (Fig. 5a). By streamlining the PCP protocol from 24 to 14 fractions, we maintained the ability to identify clusters corresponding to 11 distinct compartments (Fig. 5b-c, Extended Fig. 3a-c). The reduced number of fractions resulted in lower resolution and higher F1 values for the major compartment associations with a weighted average of 0.72 for control condition (Extended Fig. 3d). However, considering multiple predictions C-COMPASS identified 80-100% of marker proteins within their classes (Fig. 5d, Extended Fig. 3e). Endosomal, Golgi apparatus and lysosomal proteins had lower accuracy due to limited uniqueness of their profiles across gradients (Fig. 5b, Extended Fig. 3b, 3d-e). The protein distribution was like that in previous studies, with 37% distributed across multiple compartments and only 0.5% showing localization in four different compartments (Fig. 5e-h, Extended Fig. 4a-c).

**Figure 5.**
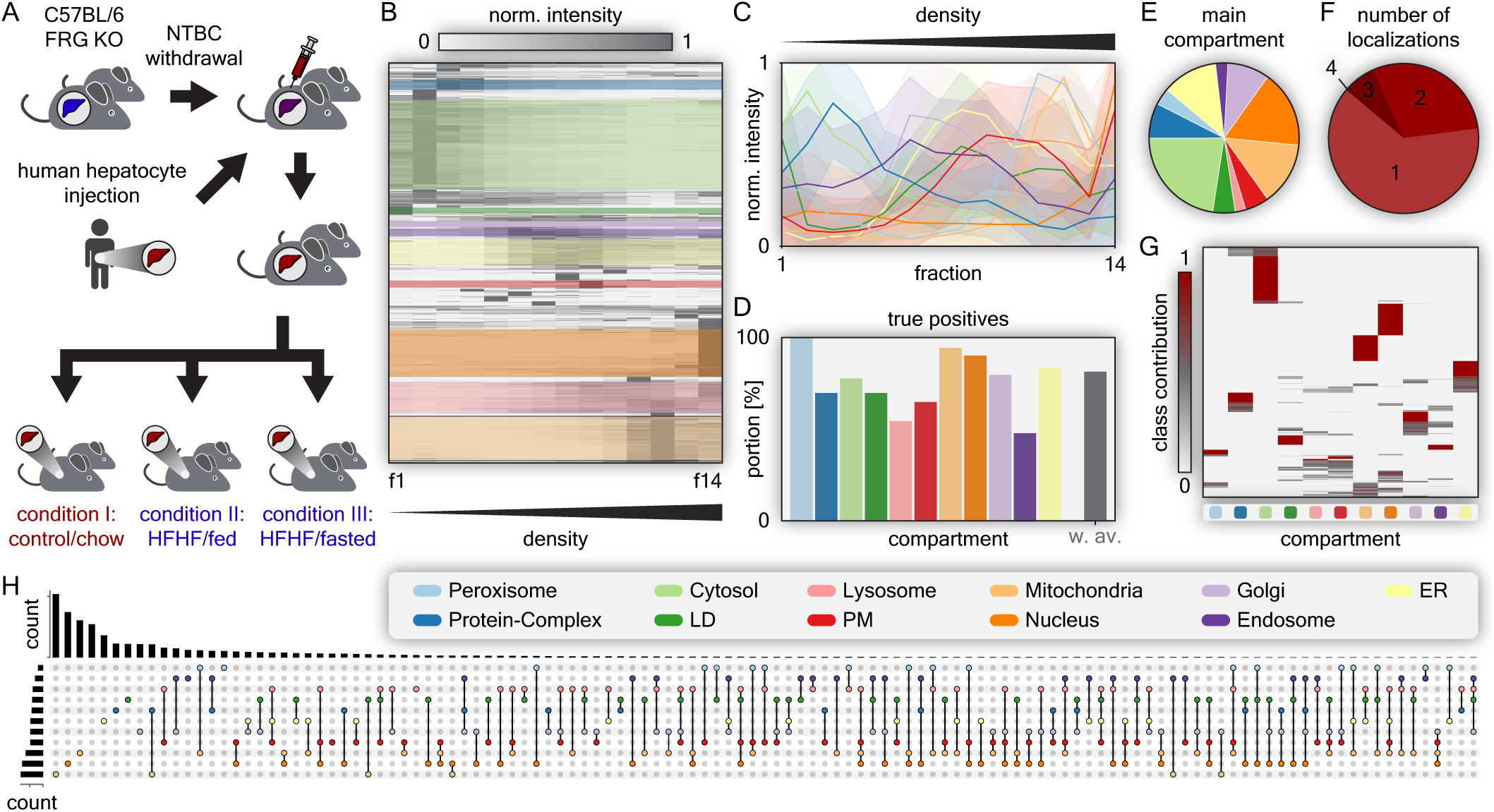
An organellar map of humanized liver. (A) Mice with humanized liver were fed either a chow diet or a HFHF diet for 5 weeks, and livers were collected in *ad libitum* fed state. A third group on the HFHF diet was subjected to an overnight fast. Livers from all three conditions were used to perform the PCP workflow. (B) Hierarchical clustering heatmap of protein profiles across 14 fractions in the chow state. Colored areas indicate clusters with the highest enrichment scores for specific compartment marker annotations. (C) Profiles of compartment markers, with line values representing median values derived from median profiles across replicates, and error bands indicating standard deviations in the chow state. (D) Bar plot showing the number of true positive localization predictions by checking for positive CC values for each class in the chow state. The grey bar represents the average true positive value weighted by batch sizes of marker proteins. (E) Number of main localization assignments to compartments in the chow state. (F) Number of proteins assigned to single or multiple compartments in the chow state. (G) Heatmap of CC values across all compartments in the chow state. (H) List of all identified compartment combinations and their frequencies in the chow state.

In our experiment, we consistently identified 7,690 proteins across three metabolic conditions. Half of these proteins maintained the same main organelle prediction when comparing chow-fed to HFHF-fed conditions, or HFHF-fed to HFHF-fasted conditions. Around a quarter of the proteins exhibited a RLS <1, indicating minor changes, while the remaining proteins were evenly split between fully relocalized (RLS = 2) and partially relocalized (RLS between 1 and 2) (Fig. 6a). Of the ∼2000 proteins with RLS >1 in both comparisons 1,300 overlaid in both conditions, indicating that the localization of this set of proteins is highly dynamic and controlled by the metabolic state (Extended Fig. 5a). All compartments were affected to varying extents, with LDs showing a particularly high percentage of relocalization events and mitochondria being underrepresented (Fig. 6b-c, Extended Fig. 5b-c). To identify candidates with the strongest and most reliable relocalizations, we further compared the C-COMPASS distance between conditions along with the corresponding p-values for proteins with a RLS >1. This analysis allowed us to pinpoint the most significant changes in protein localization (Fig. 6d, Extended Fig. 5d). One example of a protein with high RLS and high confidence is PLIN5, which localizes to LDs to mediate contact with the mitochondria. PLIN5 undergoes phosphorylation and activation during starvation^32^, facilitating fatty acid channeling for beta-oxidation and lipophagy^33,34^. Under chow condition, PLIN5 was fully localized to LDs. However, under HFHF condition, PLIN5 relocalized to the cytosolic fraction, as indicated by its increased presence in the second fraction. C-COMPASS analysis revealed a CC for PLIN5 of 63% in the cytosol and 37% on LDs. Following fasting after the HFHF diet, PLIN5 shifted back to a CC of 100% on LD (Fig. 6e). Another example of protein relocalization involves the conserved oligomeric Golgi (COG) complex, essential for organizing the Golgi apparatus^35^. Under fasting condition, we observed a previously unreported dissociation of the COG complex lobe B subcomplex. All components of this subcomplex exhibited a shift from the protein complex to cytosolic localization (Extended Fig. 6), highlighting how C-COMPASS can unveil unknown relocalization events.

**Figure 6.**
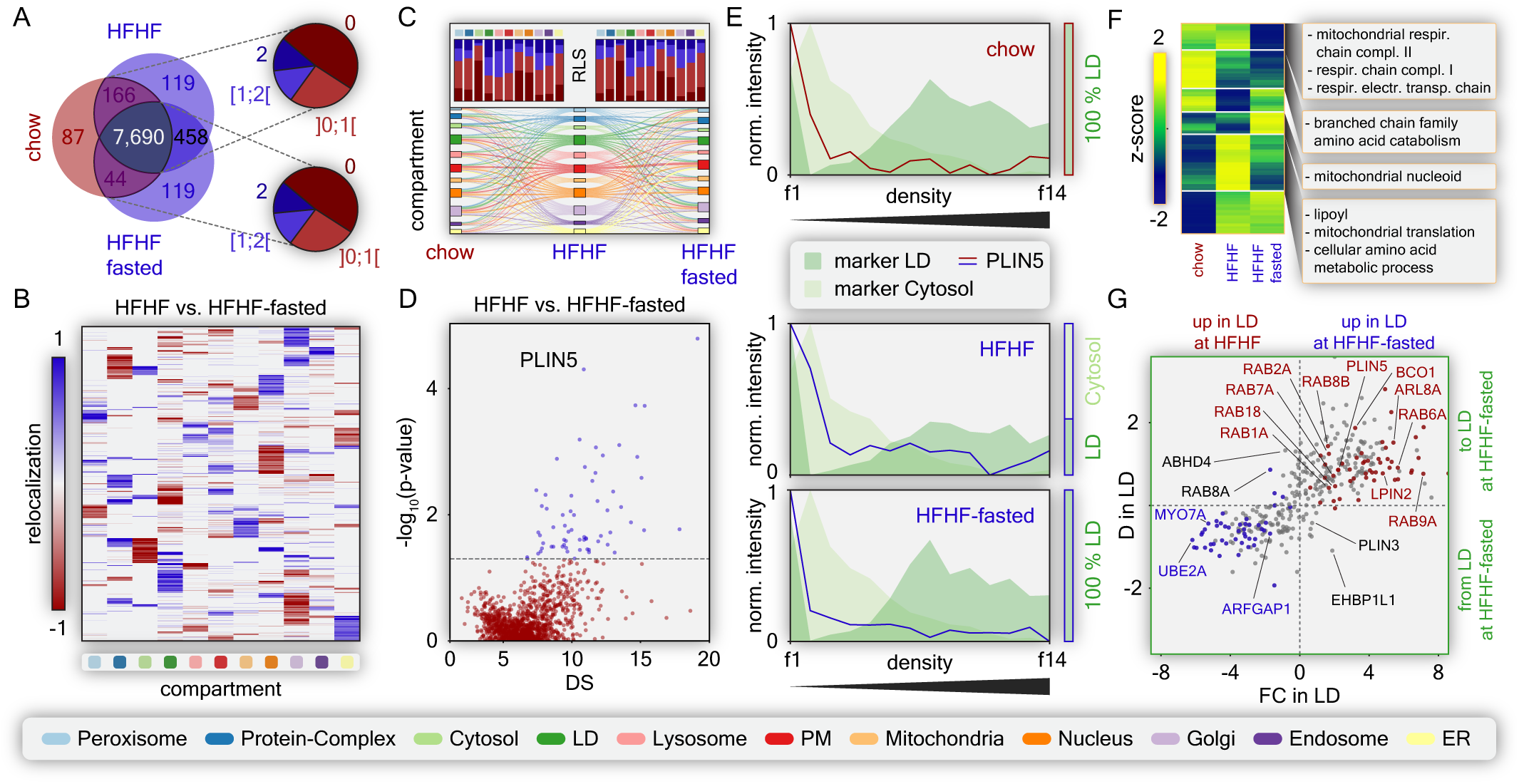
Characterization of metabolic state induced relocalization and organelle remodeling in humanized livers. (A) Venn diagram indicating proteins detected in all conditions and pie charts showing the number of proteins within specific RLS value ranges for the comparison of chow vs. HFHF (upper) and chow vs. HFHF-fasted (lower). (B) Heatmap of RL values for each compartment across all proteins, comparing HFHF with HFHF-fasted. (C) Sankey diagram illustrating localization changes from chow to HFHF to HFHF-fasted across all compartments, filtered for RLS >0 for at least one transition, while maintaining the origin and target for each relocalization. Bar charts summarize the proteins listed for one transition in the Sankey diagram, showing the number of proteins with RLS = 0 (dark red), RLS in the range of 0 to 1 (bright red), RLS between 1 and 2 (bright blue), and RLS = 2 (dark blue). (D) Scatter plot displaying p-values from a student’s t-test based on ensemble network output values against DS values for proteins with RLS >1, highlighting the most reliable outliers for the comparison of HFHF vs. HFHF-fasted. Grey dashed line indicates a p-value of 0.05. (E) Protein profiles of PLIN5 across the indicated metabolic conditions, overlaid with median marker profiles for LD and Cytosol. Bars on the right show the CC distributions for PLIN5 between these two compartments. (F) Unsupervised hierarchical clustering of proteins associated with Mitochondria, corrected for protein expression levels and organelle abundance across metabolic conditions. GO annotations, KEGG pathways, and keywords enriched in the compartments are indicated (one-sided Fisher’s exact text, FDR <0.15). (G) Scatter plot of proteome changes within LD between HFHF and HFHF-fasted. D values for Mitochondria vs. fold changes (RL values corrected by expression and abundance). Proteins with RL >0.5 are red, RL <-0.5 are blue. Red in the upper right shows higher LD localization and abundance in HFHF-fasted; blue in the lower left shows the opposite.

Finally, we applied the extended C-COMPASS pipeline to assess cellular changes in an organelle-centric manner. This approach integrates information on protein localization, protein levels, and the overall quantity of each compartment, derived from parallel total proteome analyses, to adjust protein distribution across compartments. By comparing these normalized values across different conditions, we identified proteins with changed levels in specific organelles, predicting functional changes in these organelles (Extended Fig. 7). As exemplified for mitochondria, enrichment analysis of Gene Ontology (GO) annotations, Kyoto Encyclopedia of Genes and Genomes (KEGG) pathways, and Keywords, within the distinct protein clusters, highlighted functional reprogramming under different conditions (Fig. 6f). This analysis showed metabolic changes, such as decreased respiratory capacity in the HFHF-fasted state, aligning with previous findings of reduced mitochondrial respiration during fasting^36^.

While changes in the mitochondrial proteome primarily stem from alterations in protein levels, the LD proteome is significantly influenced by relocalization events (Fig. 6c). Therefore, we employed an alternative approach to visualize fasting-induced changes in the LD proteome. By plotting the fold change in protein levels on LDs against LD-specific RL, we could assess how protein relocalization contributes to overall protein changes within this organelle (Fig. 6g). Most proteins either increased on LDs due to enhanced targeting or decreased in amount alongside reduced LD localization. For example, ARL8B, a protein known to target LDs^37,38^ and a homolog of ARL8B, which has been recently discovered to facilitate LD-mediated lipophagy^39^, exhibited increased abundance and relocalization to LDs. Conversely, the ubiquitin conjugating enzyme UBE2A was observed to relocate away from LDs, accompanied by a decrease in its LD-specific levels. There were also a few proteins that increased in abundance despite reduced targeting such as PLIN3 or EHBP1L1, or vice versa such as RAB8A or ABHD4.

### Integration of multi-omics Data with C-COMPASS

Organelle functions are defined not only by their specific proteins but also by their distinct lipid compositions. However, comprehensive organelle lipid maps have not been generated due to the scarcity of organelle-specific lipid markers. To address this gap, we evaluated the capability of C-COMPASS to integrate lipidomics data and predict the localization of lipid classes and species using neural networks trained with marker protein profiles. We conducted parallel proteomic and lipidomic analyses of organelle gradients, consisting of 14 fractions, from the livers of steatotic mice on a high-fat diet (Fig. 7a). Lipidomic analysis was performed using a quantitative, high-throughput shotgun lipidomics platform^40^, quantifying 411 individual lipid molecules across 22 lipid classes. Principal component analysis (PCA) of lipid species revealed that the primary component separating the samples was the organellar fraction, indicating distinct, organelle-specific lipid compositions (Extended Fig. 8a). This separation was validated by evaluating protein marker profiles (Fig. 7b). Comparison of lipid class profiles with protein marker profiles showed that triglycerides (TAGs) were predominantly found in the top fraction, where LDs accumulate (Fig. 7c). Cholesterol esters (CEs) and diacylglycerols (DAGs) were distributed across several fractions but were also abundant in the LD fraction. Cardiolipins (CLs), mitochondrial lipids^41^, peaked in fractions 10 to 13, corresponding to the mitochondrial protein marker peak. Phosphatidylcholine (PC) and phosphatidylethanolamine (PE), the principal phospholipids in all membranes, exhibited broad peaks across all membrane organelle fractions. PE showed an enrichment in mitochondrial fractions, confirming that the mitochondrial inner membrane is more enriched in PE compared to other cellular membranes^42^. Additionally, hierarchical clustering of individual lipid species profiles indicated a non-uniform distribution of lipid species across organelle fractions, with distinct clusters corresponding to specific organelle marker protein peaks (Fig. 7d).

**Figure 7.**
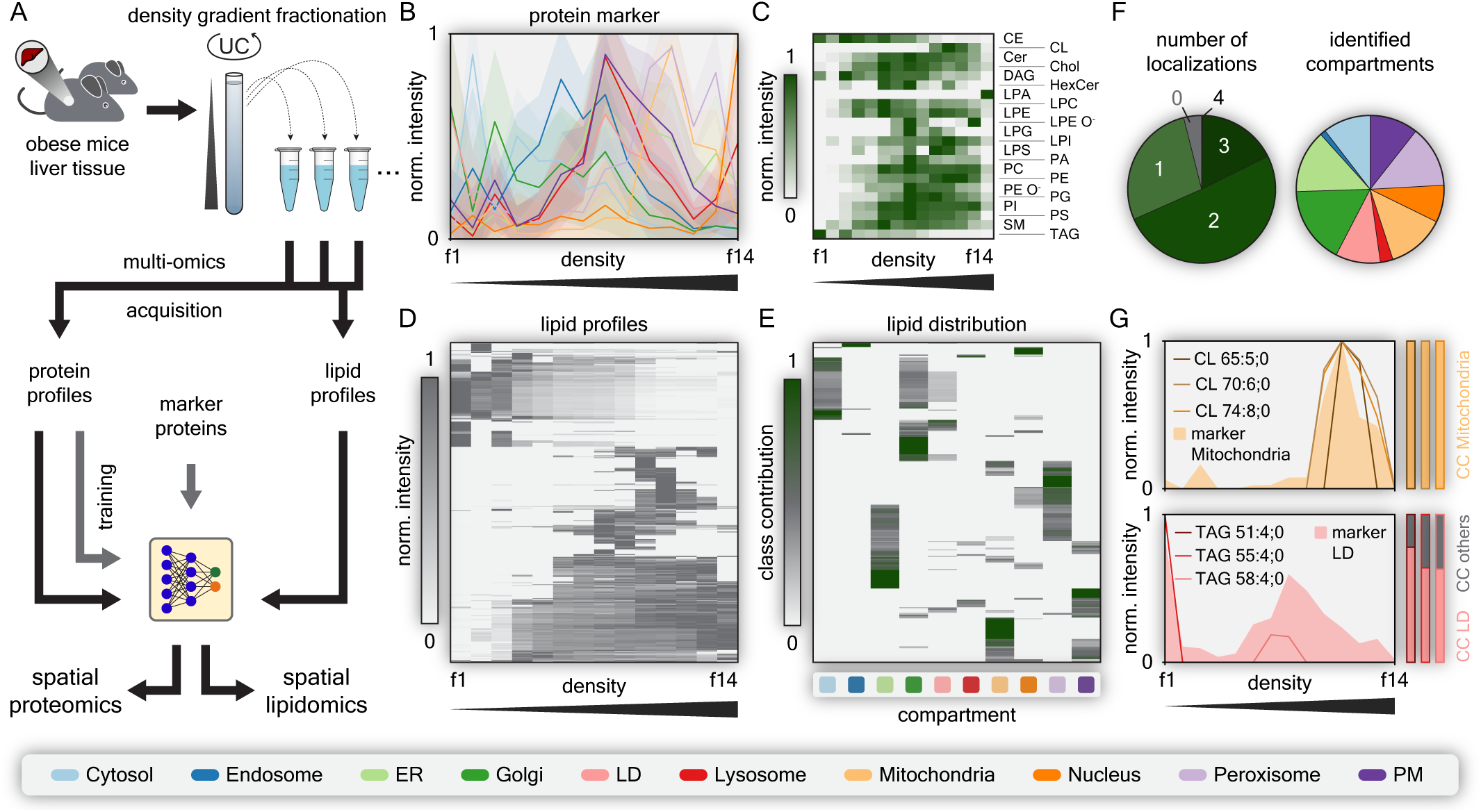
A combined protein and lipid map of steatotic liver. (A) Experimental setup scheme for integrating proteomic and lipidomic data. Liver samples from obese mice were used to perform the standard PCP workflow. All fractionation samples and total proteome samples underwent both proteomics and lipidomics analysis. After training the neural network with marker profiles derived from the proteome, both datasets were fed into the neural network for spatial predictions. (B) Profiles of compartment markers, with line values representing median values derived from median profiles across replicates, and error bands indicating standard deviations. (C) Heatmap of different lipid classes across 14 fractions. (D) Hierarchical clustering heatmap of all lipid profiles across 14 fractions. (E) Heatmap of CC values across all compartments for all identified lipids. (F) Number of lipids assigned to single or multiple compartments and the number of identified lipid localization assignments to compartments. (G) Indicated median protein marker profiles for Mitochondria (orange) and LDs (red) overlaid with distinct examples of known compartment-specific lipid species. Vertical bars indicate the percentage distribution of these lipid species across different compartments for the selected examples.

Using the C-COMPASS pipeline, we trained the neural network-based regression model with the protein marker profiles from the co-generated proteomics data and predicted lipid species compartmentalization coefficients (CC values) across organelles (Fig. 7e). Unlike proteins, where more than half exhibit unique cellular localizations, most lipids demonstrated multiple localizations. The highest lipid complexity and the greatest number of species were observed in LDs, the Golgi apparatus, mitochondria, and peroxisomes (Fig. 7f). We validated the C-COMPASS output by evaluating its organelle assignments for known organelle-specific lipid species. For instance, cardiolipin species, whose abundance profiles aligned with mitochondrial protein marker profiles, were predominantly assigned to mitochondrial localization. Similarly, triglyceride species profiles coincided with LD marker protein profiles and were largely predicted as LD-localized, although their assignment was complicated by the previously reported co-floatation of the Golgi apparatus with LDs in steatotic liver (Fig. 7g). Additionally, ceramides, synthesized in the ER and modified during their journey through the secretory pathway^43^, were primarily predicted to localize in the ER (Extended Fig. 8b). To refine our analysis, we tested an alternative approach by removing neutral storage lipids (TAG, DAG, CE) from the dataset and performing median normalization prior to profile scaling (Extended Fig. 8c-d). This method allowed us to compare phospholipid compositions of organelle membranes without the levels being confounded by substantial amounts of storage lipids in a few fractions. Specifically, we compared the phospholipid composition of LDs and the ER from which they derive. Our analysis revealed that some phospholipids were equally present in both organelles (e.g., PE 18:1/18:2), while others were more enriched in the ER (e.g., PC 15:0/18:2) (Extended Fig. 8e).

We leveraged the C-COMPASS outputs and pipeline to: (I) assess the distinct lipidomic characteristics of organelles, focusing on features such as fatty acid length and unsaturation, and (II) predict the distinct organelle lipidomes. For (I), we plotted the total number of double bonds and carbon atoms for all lipid classes across different compartments. The density plot revealed differences in the carbon chain length and unsaturation distribution between organelles. For example, lysosomes and the plasma membrane consisted of lipid species with a more defined range of carbon atoms, but higher unsaturation compared to the ER, which had a broader range of carbon atoms and more saturated lipids (Extended Fig. 8f). For (II), alongside the organelle-centric proteome analysis, we used the C-COMPASS pipeline to perform an organelle-centric lipidome analysis to predict organelle lipid compositions. However, this prediction faced limitations due to incomplete data, as not all lipid species detected in organelle fractions were present in the total liver lipidome, which is crucial for comprehensive organelle modeling (Extended Fig. 8g). This resulted in decreased modeling accuracy. Despite these limitations, our findings demonstrate that it is feasible to predict lipid localizations and that the C-COMPASS pipeline can reveal differences in membrane composition between compartments, providing insights into the distinct lipidomic profiles of cellular compartments. This approach allows for the generation of integrated cellular maps at both protein and lipid levels and can be extended to other omic layers, such as metabolomics or transcriptomics, to provide a comprehensive multi-omic view of cellular architecture.

## Discussion

In this study, we introduce C-COMPASS, an open-source software designed for analyzing proteome-wide spatial distributions across multiple cellular compartments using neural network-based regression models. C-COMPASS provides a comprehensive and user-friendly analysis pipeline with a graphical user interface, making it accessible to a broad scientific community and applicable to a wide range of biological questions across various model organisms and research fields. We validated C-COMPASS with diverse datasets, showing its ability to accurately predict multi-organelle localization and model organelle composition at both the protein and lipid levels under different biological conditions. Moreover, the pipeline can be extended to any omic-level of interest. The unique integration of lipid and protein mapping enables the creation of the first systematic organelle map at the lipid level, opening new avenues for exploring research questions.

Our results show that C-COMPASS effectively overcomes the limitations of previous methods by addressing the complex multilocalization patterns of proteins, which previous methods struggled with, allowing only single classifications^12,15^. For example, the application of C-COMPASS to a dataset of human white adipocytes demonstrates significant improvements in prediction accuracy compared to traditional SVM-based methods. The inclusion of upsampling and neural network training enhanced the precision, recall, and F1 scores across various compartments, especially for underrepresented organelles in the marker set. This improvement is evident in the increased identification of LD proteins. We further demonstrate the versatility of C-COMPASS by its applicability to datasets generated with different fractionation and labeling approaches, such as LOPIT and PCP. We show that the number of fractions used increases prediction accuracy, although there is a tradeoff between measurement time, experimental workload, and organellar resolution, which must be considered in the experimental design. Furthermore, C-COMPASS goes a step further by modeling compartment composition through integrating information on protein and organelle abundance. This allows for a comprehensive characterization of the functional evolution of compartments across biological conditions.

C-COMPASS extends its workflow to simultaneously map the protein and lipid compositions of organelles. Previously, lipid assignment based solely on lipidomics data was challenging due to a lack of organelle-specific markers. The parallel generation of protein and lipid maps now allows for the localization of lipid classes and species. This approach enables for the first time the identification of organelle-specific lipids and variations in membrane lipid composition in a comprehensive systematic manner. Traditionally, studies have focused on either proteomics or lipidomics, limiting the understanding of cellular functions and dynamics. By integrating these two approaches, we can achieve a more comprehensive view of cellular processes, revealing the interplay between proteins and lipids within cellular compartments. Proteins and lipids are the key components of cellular membranes and organelles, influencing their structure, function, and dynamics. Proteins often rely on lipid environments for localization, stability, and function, while lipids are affected by protein interactions and signaling pathways. By generating parallel protein and lipid maps, we offer a solution to investigate how specific lipid species impact protein localization and function, explore mechanisms of membrane organization, and study signaling pathways and metabolic processes mediated by protein-lipid interactions. This co-analysis offers new insights into how changes in lipid composition affect protein function and contribute to cellular dysfunctions and diseases.

While C-COMPASS provides an easily accessible solution for generating organellar maps at both protein and lipid levels for various models and biological questions, certain limitations must be considered. The software cannot overcome experimental resolution constraints; compartments with similar profiles or those that co-fractionate can result in mispredictions. For example, in steatotic liver Golgi apparatus proteins co-fractionate with LDs, as previously reported^12^, making the organelle assignment challenging. Therefore, Users should assess the uniqueness of profiles prior application of the workflow and evaluate error rates for compartments. Additional fractionation steps or incorporation of complementary fractionation procedures can improve separation but increase workload and MS time, requiring larger sample amounts. Biological replicates are crucial in *in vivo* experiments, although high workload and sample numbers can limit their use. Moreover, the method is not absolutely quantitative. It can detect quantitative changes when comparing conditions but does not measure the absolute amounts of protein in compartments. Additional experimental factors, such as protein amount per fraction and the material loaded onto MS, influence quantification. Moreover, Bayesian neural networks might be used to further quantify the prediction uncertainties and could furthermore improve the workflow.

In conclusion, C-COMPASS offers an easy and broadly applicable tool for analyzing cellular compartmentalization, addressing limitations of traditional methods such as SVMs. Its ability to predict multi-organelle localization and integrate protein and organelle abundance data provides a more comprehensive understanding of cellular dynamics. Applying C-COMPASS to various biological and disease conditions across different organs or cell models demonstrates its potential for new discoveries through systematic multi-omic organelle profiling.

## Material and Methods

### Mouse models

6-week-old female C57BL/6N FRG KO mice repopulated with human hepatocytes (#HHF13022, PNPLA3 I148M +-/-+), purchased from Yecuris, were housed at Regeneron Pharmaceuticals New York Medical College under controlled conditions (12-hour light/dark cycle, 22 ± 1 °C, 60%-70% humidity). The mice were fed *ad libitum* with either standard low protein chow (5LJ5, Purina Laboratory Rodent Research Diets Inc. 5001) or a high-fat high-fructose diet (HFHF) (D09100310iA18080901i; Research Diets Inc.) for 5 weeks. For the fasted condition, food was withdrawn overnight for 18 hours. All mice were housed within an AAALAC-accredited mouse production barrier facility at Regeneron Pharmaceuticals, Inc. New York Medical College in Valhalla Tarrytown, NY. At the age of 8 months, mice were euthanized. All procedures were approved by New York Medical College’s Regeneron Pharmaceuticals’ Institutional Animal Care and Use Committee and were conducted in accordance with the Guide for the Care and Use of Laboratory Animals, Eighth Edition, and the AVMA Guidelines for Euthanasia of Animals: 2013 or 2020 Editions.

Male C57BL/6J mice for combined lipid and protein correlation profiling, were double-housed and kept under constant ambient conditions of 22 ± 1 °C, 45–65% humidity and a 12hr/12hr light/dark cycle, with lights on from 6am until 6pm. Mice had free access to water and were fed *ad libitum* with a high-fat diet (D12331, 58% kcal fat; Research Diets, New Brunswick, NJ, USA). At the age of 43 weeks, mice were euthanized and livers perfused post-mortem with ice-cold PBS before subjecting them to organelle fractionation. Experiments were performed in accordance with the animal protection law of the European Union and upon permission from the local animal ethics committee of the Government of Upper Bavaria, Germany.

### Organelle fractionation

For each experiment, half of a mouse liver was used. Protein correlation profiling was performed with 3-4 biological replicates. PBS-perfused livers were isolated and homogenized on ice using a tissue homogenizer in 2.5 ml of buffer (20% sucrose, 20 mM Tris pH 7.4, 0.5 mM EDTA, 5 mM KCl, 3 mM MgCl_2_, protease inhibitor, and phosphatase inhibitor cocktail from Roche). A 500 µl aliquot of the lysate was reserved for total liver proteome analysis. The remaining 2 ml of lysate was centrifuged at 500 x g for 15 minutes to pellet the nuclei. The 2 ml supernatant was placed on top of a continuous 11 ml 20%-50% sucrose gradient in 20 mM Tris pH 7.4, 0.5 mM EDTA, 5 mM KCl, 3 mM MgCl_2_, with protease and phosphatase inhibitors. Subcellular organelles were separated by sucrose-density centrifugation at 100,000 x g for 3 hours at 4 °C using a Beckman SW41 Ti rotor. LDs were isolated by collecting the 1 ml top fraction with a tube-slicer (Beckman Coulter). The remaining 1 ml gradient fractions were collected sequentially with a pipette from top to bottom. In total, 14 fractions were collected from each biological replicate and condition for proteomic analysis.

## Proteomic analysis

### Proteomic sample preparation

Protein concentration of samples was measured using the BCA Protein Assay (Thermo, 23225). 50 µg of protein was adjusted (2% sodium deoxycholate and 100 mM Tris-HCl, pH 8.5) and boiled for 5 minutes at 95 °C and 1,000 rpm. Samples were sonicated using a Diagenode Bioruptor for 15 cycles of 30 seconds. Proteins were digested overnight at 37 °C and 1,000 rpm with trypsin (Sigma, t6567) and LysC (Wako, 129-02541) at a 1:50 protein-to-enzyme ratio. Proteins were reduced and alkylated with 10 mM tris(2-carboxyethyl)phosphine (TCEP) and 40 mM chloroacetamide at 40 °C and 1,000 rpm in the dark for 10 minutes. Peptides were then acidified by adding an equal volume of isopropanol and 2% trifluoroacetic acid (TFA). After centrifugation at 15,000 x g for 10 minutes, supernatants were loaded onto activated triple-layer styrenedivinylbenzene reversed-phase sulfonated Stage Tips (3M Empore). The peptides were washed sequentially with 100 µl ethyl acetate containing 1% TFA, 100 µl 30% methanol containing 1% TFA, and 150 µl 0.2% TFA, and then eluted with 60 µl elution buffer (80% acetonitrile and 5% ammonium hydroxide). Peptides were lyophilized and dissolved in 10 µl MS loading buffer (2% acetonitrile and 0.1% TFA).

### Proteomic Data Acquisition

For LC-MS/MS analysis, 500 ng of peptides were analyzed using an Orbitrap Exploris 480 (Thermo Fisher Scientific) equipped with a nano-electrospray ion source and FAIMS (Thermo Fisher Scientific). Fractionation samples were subjected to a single CV at -50 V, while total proteome samples used dual CVs at -50 and -70 V. The system was coupled with an EASY-nLC 1200 HPLC (Thermo Fisher Scientific). Separation of peptides occurred at 60 °C on a 50 cm column with 75 µm inner diameter, packed in-house with ReproSil-Pur C18-AQ 1.9 µm resin (Dr. Maisch). The gradient was set to 60 minutes for fractionation samples and 115 minutes for total proteome samples, employing reversed-phase chromatography with a binary buffer system: buffer A (0.1% formic acid) and buffer B (80% acetonitrile, 0.1% formic acid). Initially, buffer B startet at 5% and increased to 45% over 45 minutes for fractionation samples and 95 minutes for total proteome samples, followed by a washout phase at 95%, all at a flow rate of 300 nl/min. Peptides were ionized via electrospray ionization and transferred to the gas phase. Data acquisition utilized a DIA tandem mass spectrometry method with variable window sizes. Each cycle included one MS1 scan (300 – 1,650 m/z, maximum ion fill time of 45 ms, normalized AGC target of 300%, R = 120,000 at 200 m/z), followed by fragment scans of 66 unequally spaced windows for humanized liver fractionation samples and 33 windows for high-fat diet mouse liver fractionation samples and all total proteome samples (fill time of 22 ms, normalized AGC target of 1,000%, normalized HCD collision energy of 30%, R = 15,000). Spectra were recorded in profile mode with positive polarity.

### Proteomic Data Processing

DIA raw data files were processed using Spectronaut 18 (Copernicus, Biognosys) with the directDIA+ deep-mode option, searching against the Uniprot databases for human (UP000005640_9606) and mouse (UP000000589_10090) for liver samples. Trypsin/P cleavage allowed peptide lengths of 7-52 amino acids with up to two missed cleavages. Fixed modifications included carbamidomethylation, and variable modifications included methionine oxidation and N-terminal acetylation. The analysis selected 3-6 Best N Fragment ions per peptide with a precursor and protein q-value cutoff of 1%. Global normalization using median quantities corrected for MS intensity drift.

## Lipidomic analysis

### Lipid extraction for mass spectrometry lipidomics

Mass spectrometry-based lipid analysis was performed by Lipotype GmbH (Dresden, Germany) as described^45^. Lipids were extracted using a chloroform/methanol procedure^46^. Samples were spiked with internal lipid standard mixture containing: cardiolipin 14:0/14:0/14:0/14:0 (CL), ceramide 18:1;2/17:0 (Cer), diacylglycerol 17:0/17:0 (DAG), hexosylceramide 18:1;2/12:0 (HexCer), lyso-phosphatidate 17:0 (LPA), lyso-phosphatidylcholine 12:0 (LPC), lyso-phosphatidylethanolamine 17:1 (LPE), lyso-phosphatidylglycerol 17:1 (LPG), lyso-phosphatidylinositol 17:1 (LPI), lyso-phosphatidylserine 17:1 (LPS), phosphatidate 17:0/17:0 (PA), phosphatidylcholine 15:0/18:1 D7 (PC), phosphatidylethanolamine 17:0/17:0 (PE), phosphatidylglycerol 17:0/17:0 (PG), phosphatidylinositol 16:0/16:0 (PI), phosphatidylserine 17:0/17:0 (PS), cholesterol ester 16:0 D7 (CE), sphingomyelin 18:1;2/12:0;0 (SM), triacylglycerol 17:0/17:0/17:0 (TAG) and cholesterol D6 (Chol). After extraction, the organic phase was transferred to an infusion plate and dried in a speed vacuum concentrator. The dry extract was re-suspended in 7.5 mM ammonium formiate in chloroform/methanol/propanol (1:2:4; V:V:V). All liquid handling steps were performed using Hamilton Robotics STARlet robotic platform with the Anti Droplet Control feature for organic solvents pipetting.

### MS data acquisition

Samples were analyzed by direct infusion on a QExactive mass spectrometer (Thermo Fisher Scientific) equipped with a TriVersa NanoMate ion source (Advion Biosciences). Samples were analyzed in both positive and negative ion modes with a resolution of R_m/z=200_ = 280,000 for MS and R_m/z=200_ = 17,500 for MS/MS experiments, in a single acquisition. MS/MS was triggered by an inclusion list encompassing corresponding MS mass ranges scanned in 1 Da increments^40^. Both MS and MS/MS data were combined to monitor CE, Chol, DAG and TAG ions as ammonium adducts; LPC, LPC O^-^, PC and PC O^-^ as formiate adducts; and CL, LPS, PA, PE, PE O^-^, PG, PI and PS as deprotonated anions. MS only was used to monitor LPA, LPE, LPE O^-^, LPG and LPI as deprotonated anions, and Cer, HexCer and SM as formiate adducts.

### MS raw data analysis and post-processing

Data were analyzed with in-house developed lipid identification software based on LipidXplorer^47,48^. Data post-processing and normalization were performed using an in-house developed data management system. Only lipid identifications with a signal-to-noise ratio >5, and a signal intensity 5-fold higher than in corresponding blank samples were considered for further data analysis.

### Software for image generation

All figures were created using Adobe Illustrator version 24.3. Plots were generated in Python version 3.8.8, utilizing the following packages: NumPy (1.20.1), pandas (1.2.4), seaborn (0.11.1), Matplotlib (3.3.4), umap-learn (0.5.3), adjustText (0.8), SciPy (1.6.2), scikit-learn (0.24.1), UpSetPlot (0.9.0), Plotly (5.22.0), and the Python standard libraries collections and webbrowser. Heatmaps were also created with Perseus 1.6.1.5.0.

### C-COMPASS

C-COMPASS is implemented in Python version 3.8.8 and is based on a user-centered designed paradigm. The graphical user interface of C-COMPASS was developed using PySimpleGUI (4.55.1) in combination with Tkinter (8.6). Quality metrics were calculated using scikit-learn (0.24.1), while neural network predictions were executed using Keras (2.10.0) alongside TensorFlow (2.10.0) and KerasTuner (1.4.5) for hyperparameter tuning. Additionally, C-COMPASS utilizes several libraries including NumPy (1.20.1), SciPy (1.6.2), pandas (1.2.4), and the Python standard libraries math, random, copy, os, DateTime, and collections.

### Proteomic data processing

Fractionation and total abundance data were imported into C-COMPASS, with samples assigned to conditions, replicates, and fractions. Fractionation datasets were filtered to include proteins identified in at least two replicates per condition, with missing values replaced by zeros and profiles scaled to an area under the curve of 1. Total proteome data retained proteins with at least two valid values per condition, with missing values imputed using a normal distribution (shifted by 1.8, width 0.3). Mean intensity values were used for analysis. Median profiles of replicates per condition were rescaled to a 0-1 range for visualization.

### Lipidomic data processing

Only lipids with amounts >1 pmol and identified in at least two out of three biological replicates were reported. Molar amount values (in pmol) of individual lipid (sub-)species were normalized to total lipid content per sample, yielding molar fraction values, expressed in mol%. Median profiles across replicates were rescaled to a 0-1 range for visualization. For spatial lipidomics analysis, these normalized values were scaled to an area under the curve of 1 across fractions and missing values replaced by zero. For total liver lipidome, missing values were imputed using a normal distribution (shifted by 1.8, width 0.3). Mean intensity values were used for further analysis.

### Upsampling

The batch size for upsampling was set based on the compartment with the most marker proteins, establishing this as the target. Underrepresented marker batches were supplemented with artificial profiles, generated by selecting three random marker profiles, calculating their median, adding random noise (sigma twice the standard deviation of each fraction value), and re-scaling. Profiles for two compartments were created by combining and re-scaling random profiles from each compartment. This process was repeated to generate mixed profile batches with 25:75, 50:50, and 75:25 ratios. For each combination and step, the batch size was 5% of the target for single compartments.

### Neural network architecture

Each biological replicate from the organelle separation workflow was processed independently through the neural network. The input layer size was determined by the number of fractions in each replicate dataset. This was followed by two hidden dense layers: the second dense layer had a fixed number of neurons equal to the number of classes, while the first dense layer’s size was optimized during hyperparameter tuning. This size ranged between the number of classes plus 40% to 60% of the difference between the input and second dense layer size, with optimization steps of 2. The first dense layer employed a ReLU activation function, whereas the second dense layer utilized a linear activation function. Following these, a ReLU layer was applied to produce only positive values, and a subsequent normalization layer adjusted the values so that they summed to 1 across all neurons. The output of each neuron was then used to calculate the class contribution values.

### Neural network optimization

The network optimization was conducted using a random split of the upsampled data, with 80% for training and 20% for validation. The optimization goal was to minimize the mean squared error. To prevent overfitting, an early stopping callback was employed, monitoring validation loss with a patience of 5 epochs. Hyperparameters, such as the size of the dense layer, the choice of optimizer (Adam, RMSprop, or SGD), and the learning rate (ranging from 10^-4^ to 10^-1^ with logarithmic sampling), were tuned using Keras Hyperband. This tuning process was capped at 20 epochs and a factor of 3.

The entire optimization procedure – comprising upsampling, profile mixing, training, and hyperparameter tuning – was executed in three rounds for each replicate dataset, aiming to identify the best neural network architecture. Once identified, the neural network was then trained and applied to the entire dataset, including profiles with unknown localizations. This process was repeated ten times to generate an ensemble results matrix. Final prediction values were derived by averaging the output values across all rounds, followed by a re-normalization step to ensure that class values summed to 1.

### Output filtering

CC values were calculated by averaging the neural network raw outputs from the ensemble and filtering them with a threshold covering 95% of the false positive values for each compartment. Values below this threshold were set to 0, and the prediction values across all compartments were re-normalized to sum to 1 per protein. Proteins that retained a single value greater than 1 after filtering were defined as having a single localization.

## Statistics

### Quality assessment

Quality scores, including F1, precision, and recall, were calculated using weighted averaging from scikit-learn. Each prediction was assigned to a distinct compartment based on its main localization, which is defined as the compartment with the highest CC, or (before filtering) with the highest neural network output.

### Static protein localizations

The CC values were used for static localization statistics, and the difference in CC values across conditions yielded the RL values and their p-values based on student’s t-test. Raw neural network outputs were used to calculate p-values using a student’s t-test and Cohen’s distance (D) per compartment, which measures the effect size and indicates the magnitude of differences between groups. The overall score DS was calculated by summing the absolute D values, while the RLS was determined by summing the absolute RL values derived from the processed CC data.

### Organelle composition

Compartment abundance was calculated using all proteins with the highest CC for a specific compartment, averaging their values from the total proteome. For class-centric statistics, protein intensities derived from total proteomes were multiplied by the RL values per condition and divided by the class abundance. These values were log_2_-transformed and compared between conditions to calculate class-centric fold changes. For class-centric clustering analysis, z-scoring was applied to the dataset.

## Data availability

All proteomic raw data will be made accessible on PRIDE upon manuscript publication. C-COMPASS software and software documentation will be made openly accessible upon publication. For prior use contact

## Acknowledgements

We thank Felix Klingelhuber, Maximilian Gerwien, Pamela Kakimoto and all members of the Krahmer and Hasenauer labs for discussion. We thank Daniel Brandt and Sara Ribicic for technical assistance. These studies were supported by DFG Emmy Noether (KR5166/2 to N.K.), DFG BATenergy (TRR 333/1 – 450149205 to N.K. and J.H.), DFG Germany’s Excellence Strategy (390685813 – EXC 2047 and 390 873048 – EXC 2151 for J.H.), the European Foundation for the Study of Diabetes (Future Leader Award NNF20SA0066171 to N.K.), and the University of Bonn via the Schlegel professorship (to J.H.).

## Contributions

N.K. and D.T.H. conceived the project. N.K., D.T.H and X.C. designed experiments. D.T.H. and X.M. performed organelle fractionations. E.T. and D.T.H. performed proteomic sample preparation. D.T.H. conducted proteomic analyses. D.T.H., J.H. and N.K. designed the software structure as well as the analysis and validation strategies. D.T.H. implemented software. N.K. and D.T.H. analyzed data. C.K. and M.J.G. performed lipidomic analysis and data analysis. N.K. and D.T.H. wrote the manuscript.

---

**Extended Figure 1.**
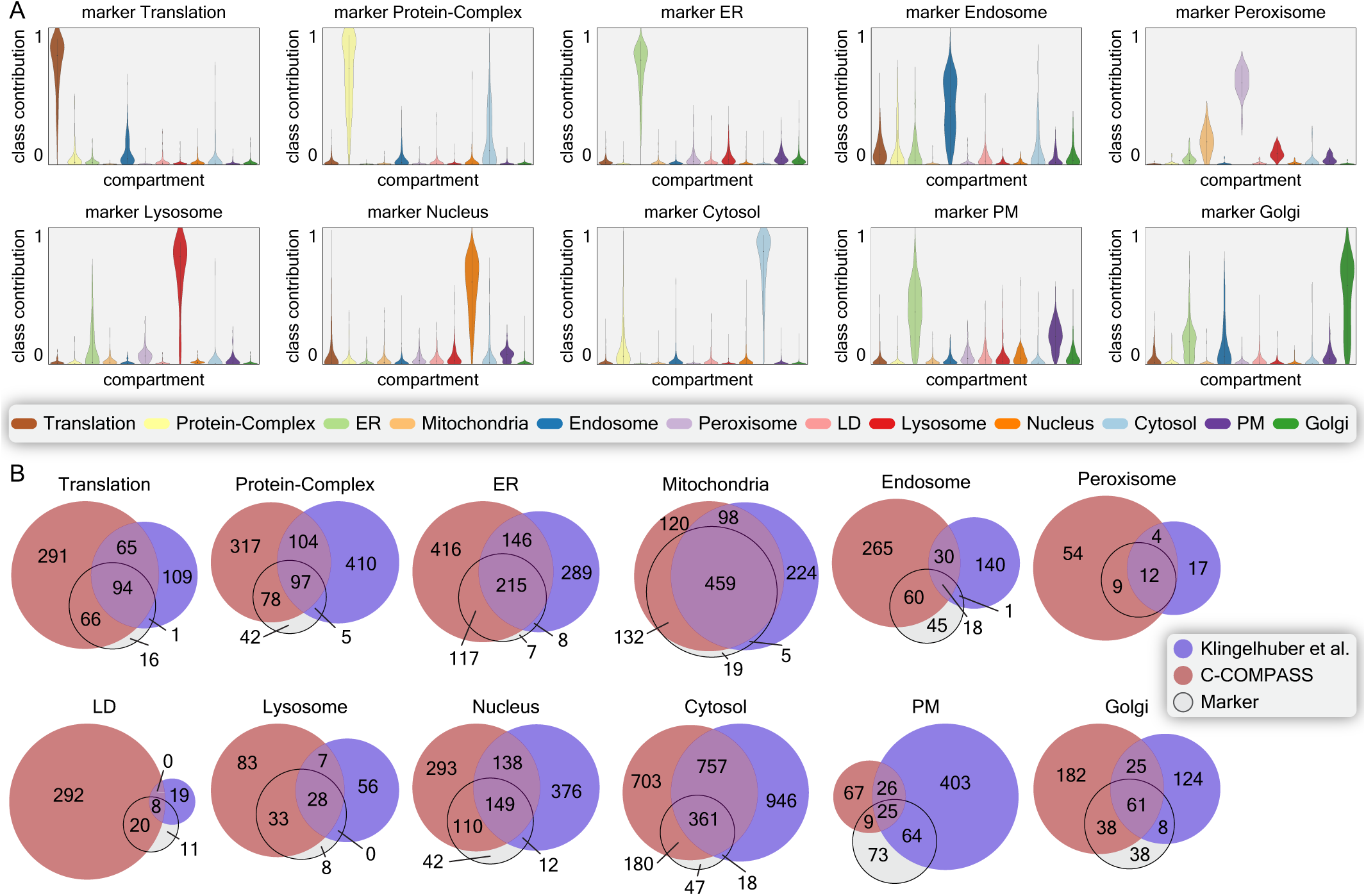
Comparison of C-COMPASS performance versus SVM predictions. (A) Violin plots show raw neural network outputs for different compartments derived from their marker proteins. Violins are width-scaled, inner lines represent quartiles, and dots represent median values. (B) Venn diagrams comparing proteins with the highest neural network output values for different compartments from C-COMPASS (red) with original predictions by Klingelhuber et al.^16^ (blue). Colorless circles show intersections with marker proteins.

**Extended Figure 2.**
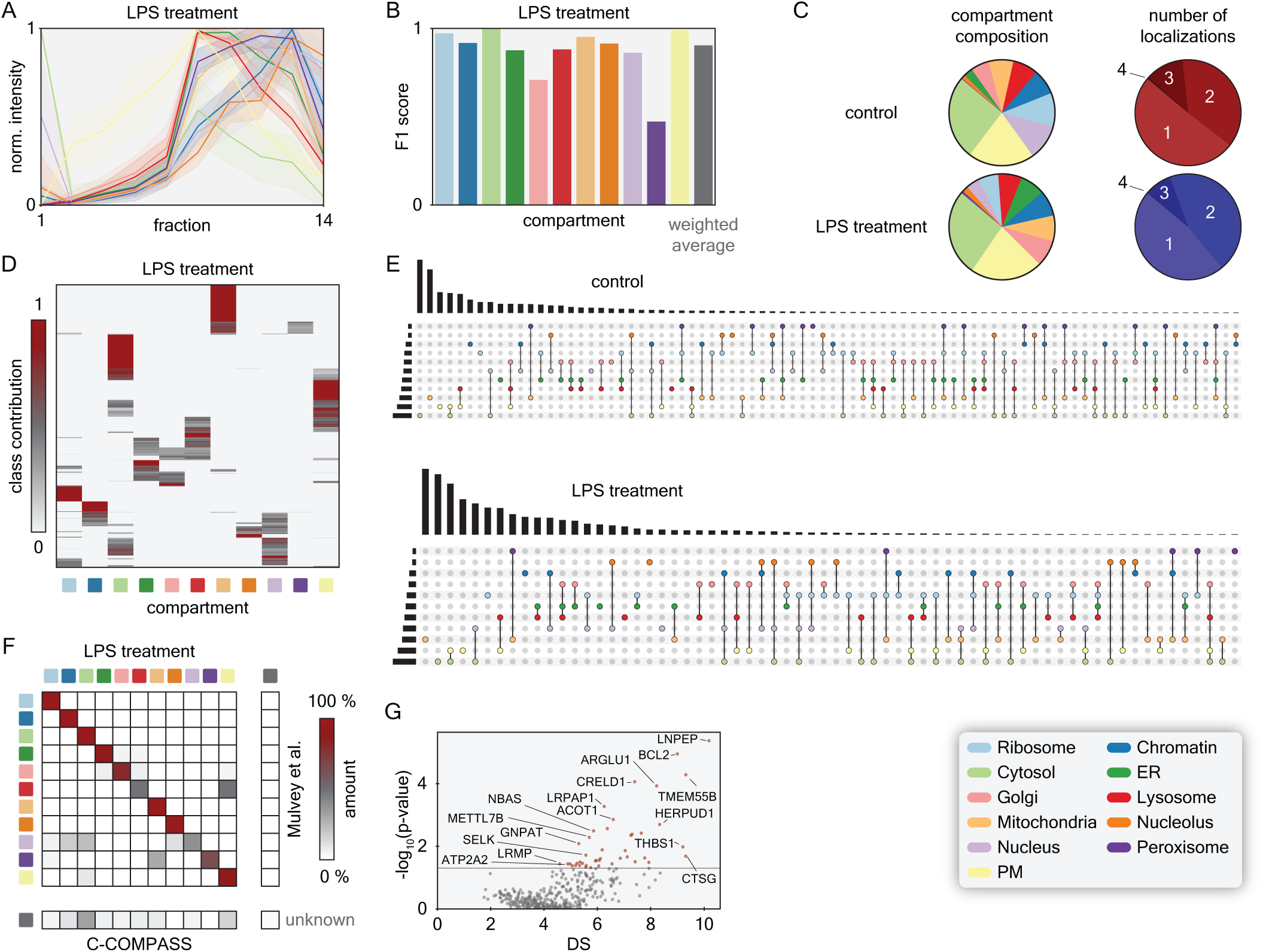
Application of C-COMPASS to the HyperLOPIT workflow. (A) Profiles for compartment markers, with line values representing median values derived from median profiles across replicates for LPS treatment, and error band indicating standard deviations. (B) Bar plot showing F1 scores for each compartment for LPS treatment. Main organelle assignments were determined using the highest CC value per protein. The grey bar represents the average F1 value weighted by batch sizes of marker proteins. (C) Number of main localization assignments to compartments, and numbers of proteins assigned to single or multiple compartments for both conditions. (D) Heatmap of CC values across all compartments for LPS treatment. (E) Lists of all identified compartment combinations and their frequencies for both conditions. (F) Correlation matrix comparing original comparment associations by Mulvey et al.^22^ with C-COMPASS main compartment predictions for LPS treatment. (G) Scatter plot displaying p-values from a student’s t-test based on ensemble network output values against DS values for proteins with RLS >1, highlighting the most reliable outliers for the comparison between control and LPS treatment. The grey dashed line indicates a p-value of 0.05.

**Extended Figure 3.**
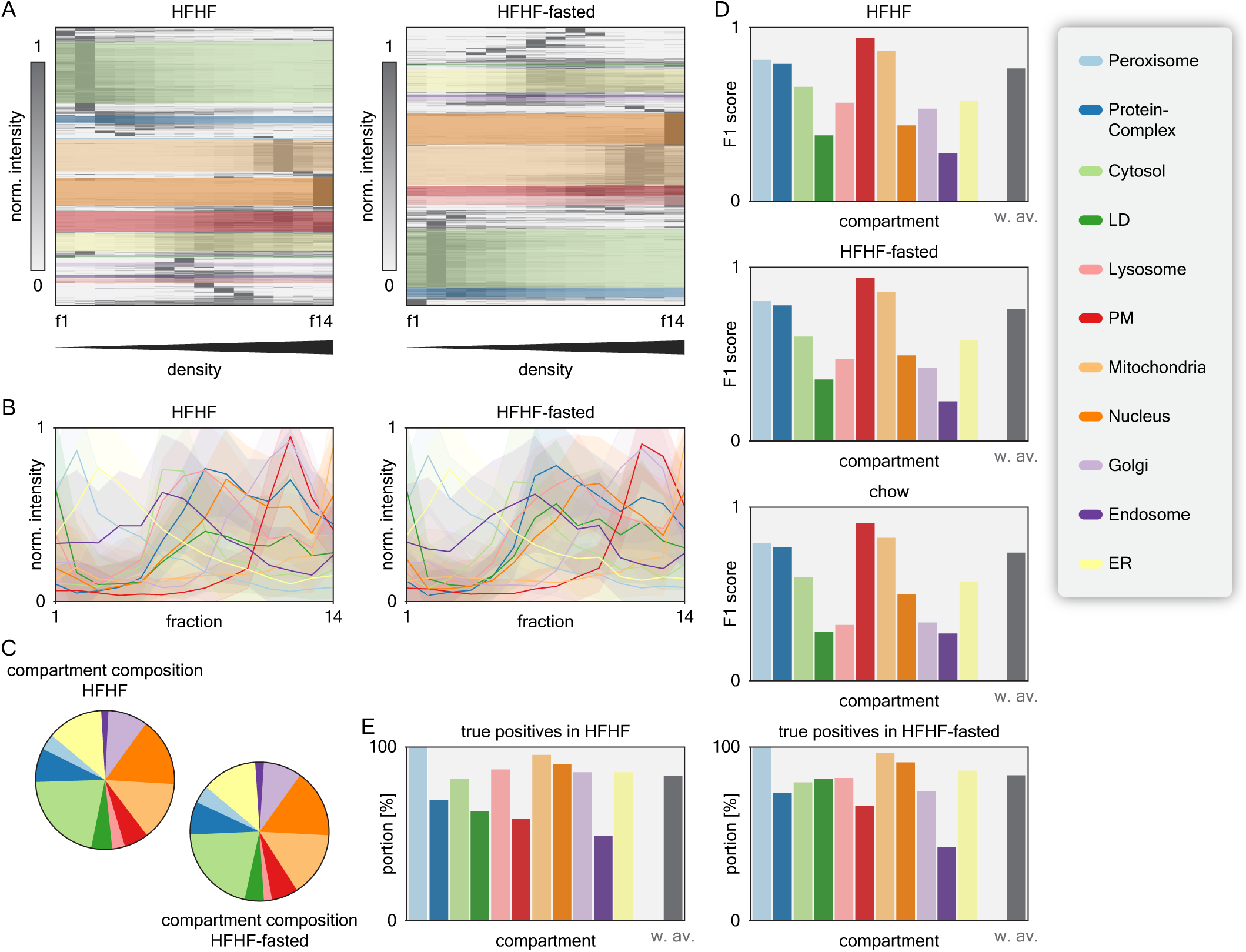
An organelle map of humanized liver across metabolic conditions. (A) Hierarchical clustering heatmap of protein profiles across 14 fractions for HFHF and HFHF-fasted conditions. Colored areas indicate clusters with the highest enrichment scores for specific compartment marker annotations. (B) Profiles of compartment markers, with line values representing median values derived from median profiles across replicates, and error bands indicating standard deviations for HFHF and HFHF-fasted conditions. (C) Numbers of main localization assignments for HFHF and HFHF-fasted conditions. (D) Bar plots showing F1 scores for each compartment across all conditions. Main organelle assignments were determined using the highest CC value per protein. The grey bar represents the average F1 value weighted by batch sizes of marker proteins. (E) Bar plot showing the number of true positive localization predictions by checking for positive CC values for each compartment for HFHF and HFHF-fasted conditions. The grey bar represents the average true positive value weighted by batch sizes of marker proteins.

**Extended Figure 4.**
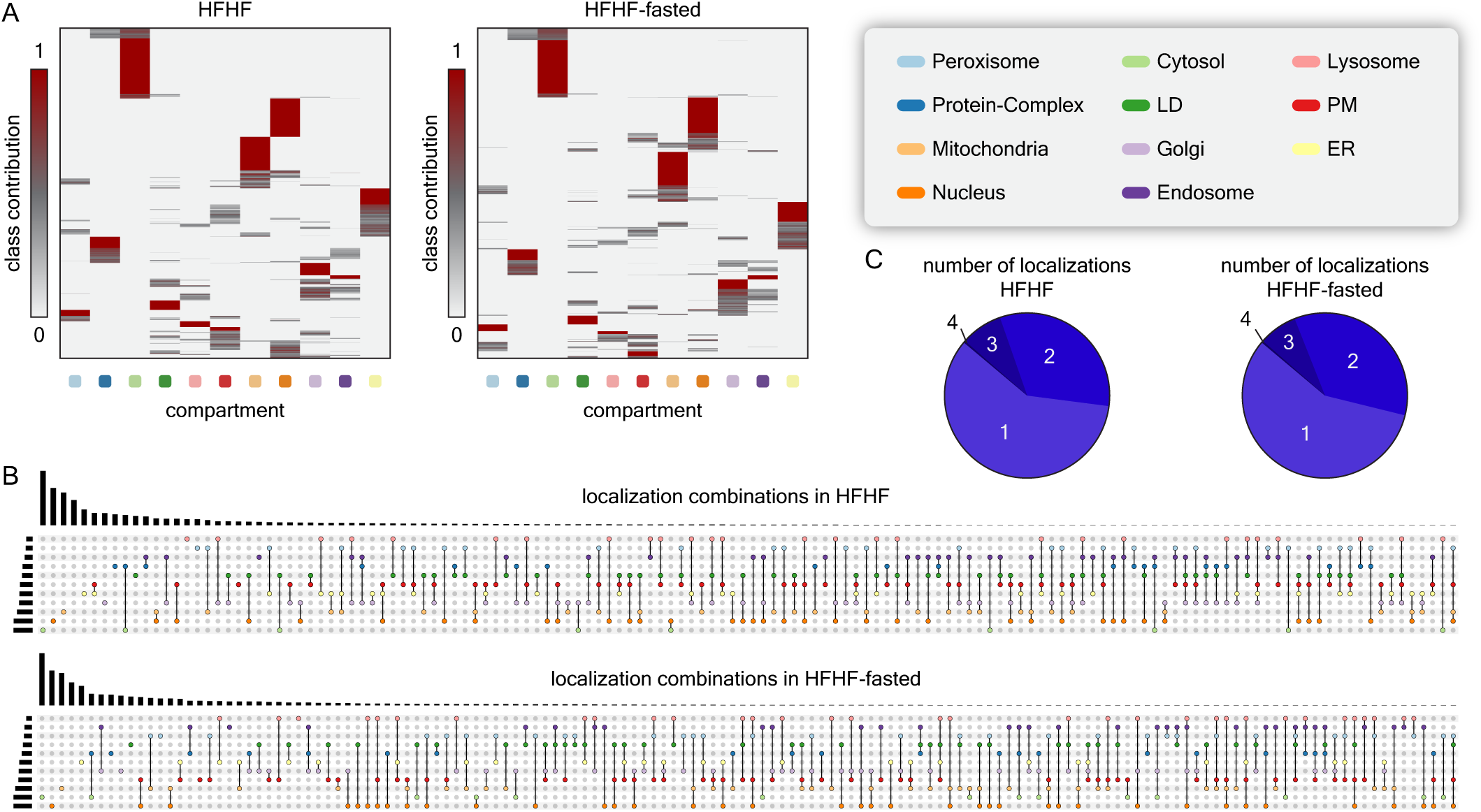
Compartment association in organelle map of humanized liver. (A) Heatmaps of CC values across all compartments for HFHF and HFHF-fasted conditions. (B) Lists of all identified compartment combinations and their frequencies for HFHF and HFHF-fasted conditions. (C) Numbers of proteins assigned to single or multiple compartments for HFHF and HFHF-fasted conditions.

**Extended Figure 5.**
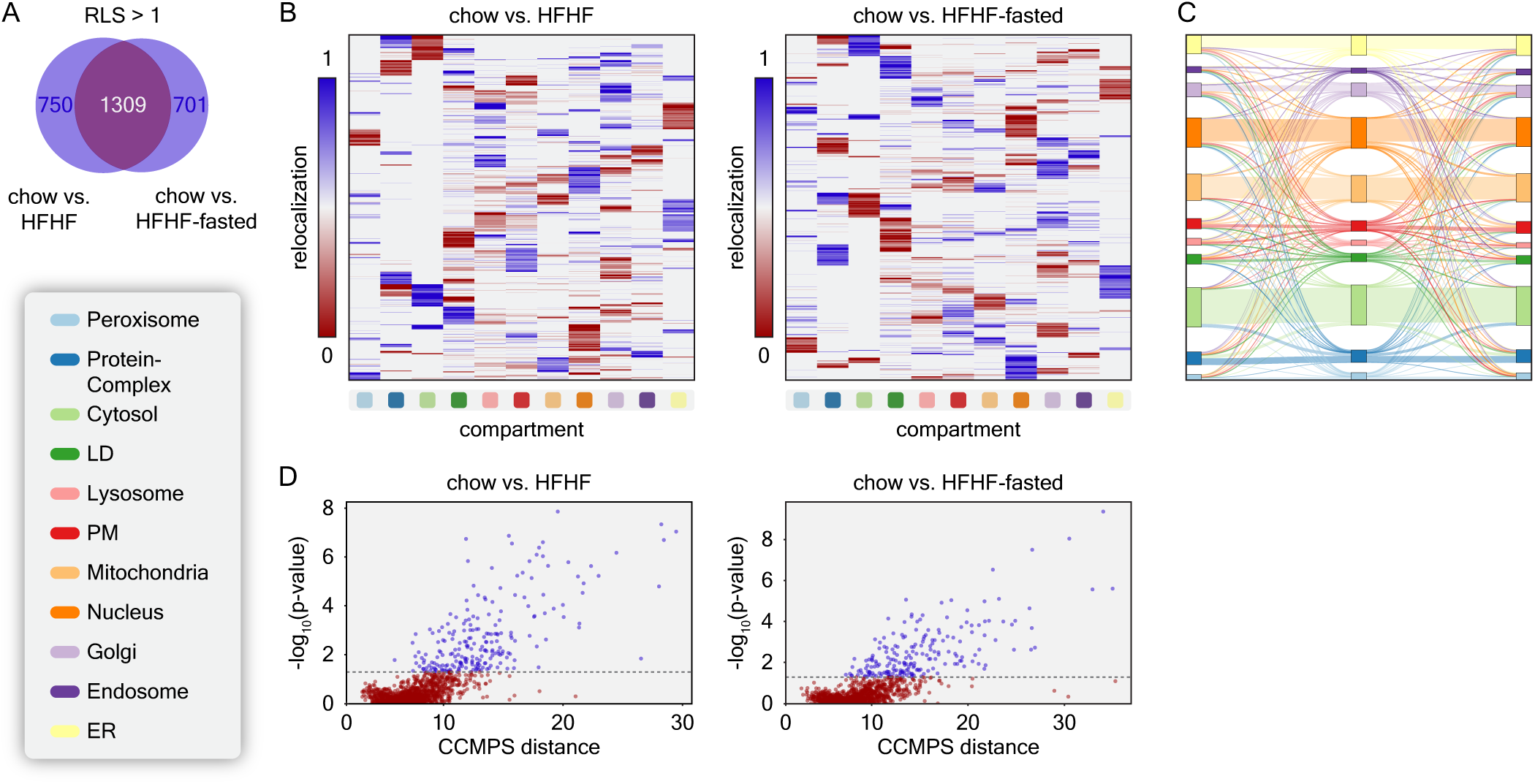
Protein relocalizations and compartmental changes across different metabolic conditions. (A) Venn diagram showing proteins with RLS >1 for the comparisons chow vs. HFHF and chow vs. HFHF-fasted. (B) Heatmaps of RL values for each compartment across all proteins for the comparisons chow vs. HFHF and chow vs. HFHF-fasted. (C) Sankey diagram illustrating localization changes from chow to HFHF to HFHF-fasted across all compartments, including non-relocalizing proteins, maintaining the origin and target for each relocalization. (D) Scatter plots displaying p-values from student’s t-tests based on ensemble network output values against DS values for proteins with RLS >1, highlighting the most reliable outliers for the comparisons chow vs. HFHF and chow vs. HFHF-fasted.

**Extended Figure 6.**
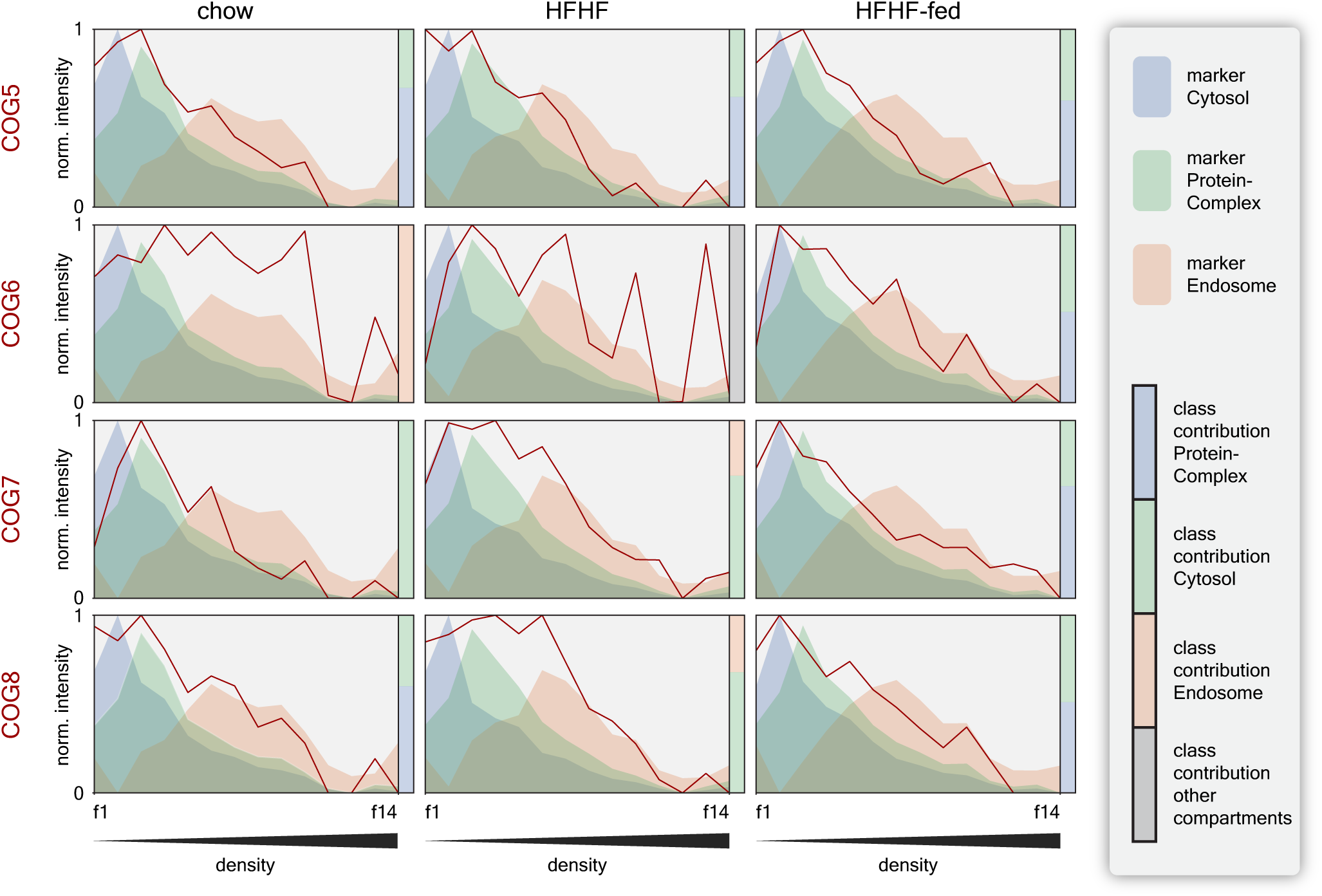
Metabolic state induced spatial re-organization of COG complex. Line plots are showing MinMax scaled median marker profiles as areas under the curve, and the distinct protein profiles derived from different COG proteins over all conditions. Bars on the right of each plot are representing the spatial distributions.

**Extended Figure 7.**
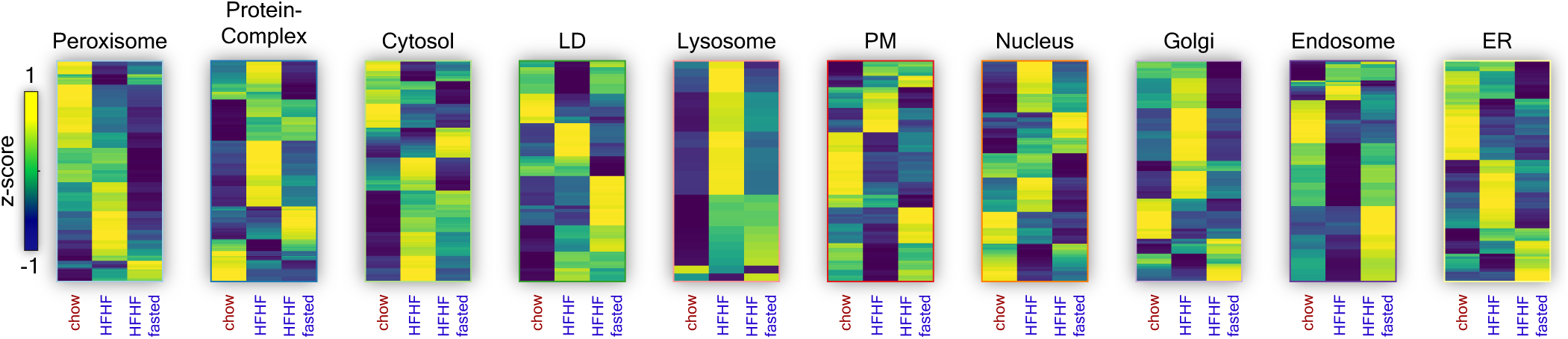
Compartmental composition changes due to metabolic states. Unsupervised hierarchical clustering of proteins associated with different compartments, corrected for protein expression levels and organelle abundances.

**Extended Figure 8.**
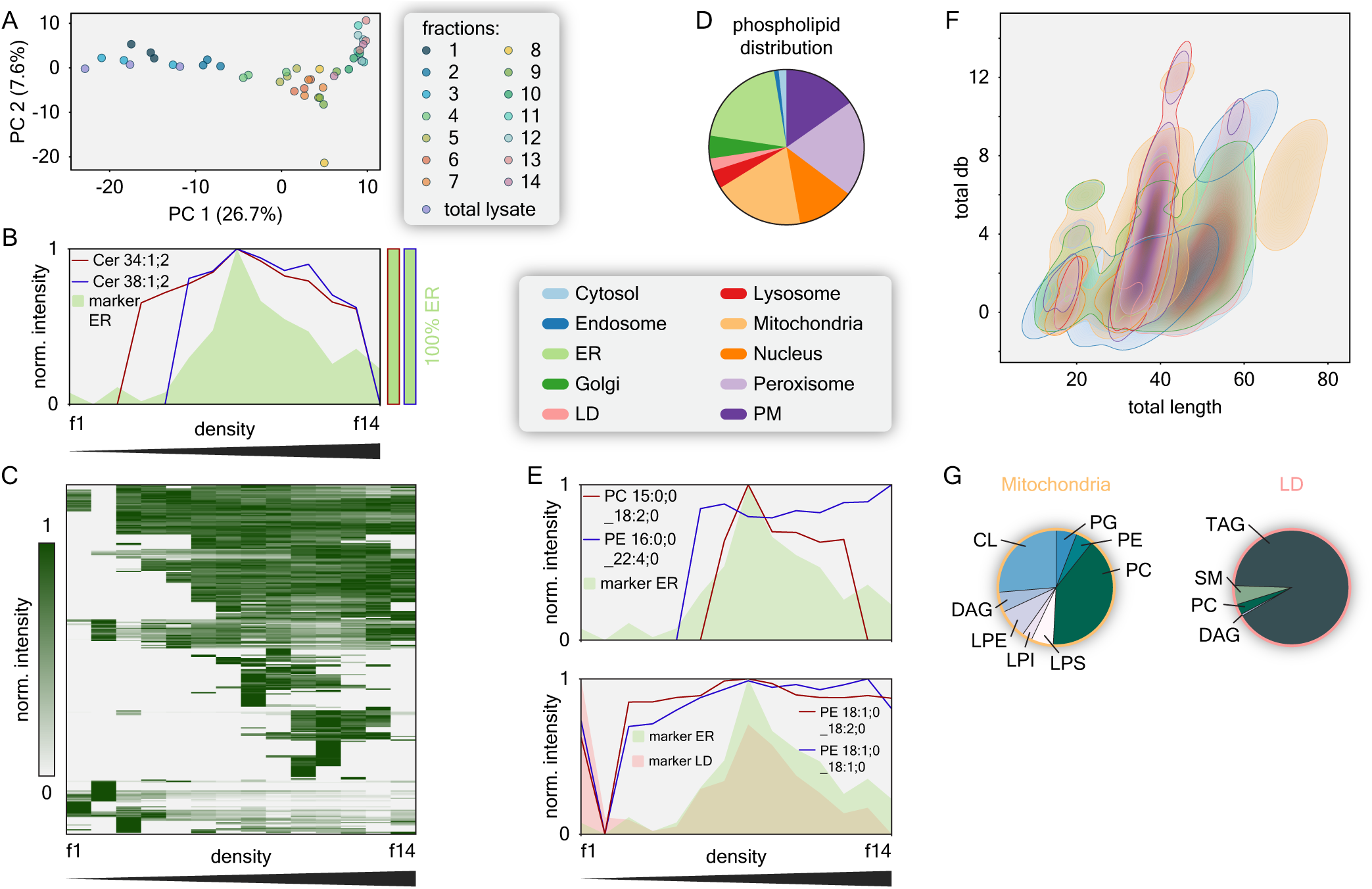
Modeling of organelle lipid composition and analysis of compartment specific phospholipid compositions. (A) Principal Component Analysis of all measured lipidomics samples. (B) Median protein marker profiles for the ER (green area) overlaid with distinct lipid profiles. Bars on the right are representing the spatial distributions. (C) Hierarchical clustering heatmap of phospholipid profiles across 14 fractions. The displayed profiles are re-normalized by width adjustment after the removal of neutral lipids. (D) Predicted phospholipid localization distribution across compartments, excluding neutral lipids. (E) Median protein marker profiles for ER (green area) and LDs (red area) overlaid with distinct phospholipid profiles. Values were re-normalized after removing neutral lipids. (F) Density plot of all lipid characteristics based on their occurrence in specific compartments (indicated by area colors). (G) Lipid class composition for Mitochondria and LDs based on values normalized to the intensities in the total lysate, as well as the relative abundance of compartments.

